# Noradrenergic Modulation of an Amygdalo-thalamic Circuit

**DOI:** 10.1101/2025.10.14.682083

**Authors:** Tracy L. Yang, Justin Bucalo, Mark L. Andermann, Chinfei Chen

## Abstract

Emotional and cognitive processing rely on communication between the basolateral amygdala (BLA) and the medial prefrontal cortex (mPFC). The BLA regulates mPFC both directly and indirectly via the medial sub-division of the medial dorsal thalamus (MDm). Although the BLA projection to MDm has been established anatomically, less is known about the functional properties of this synapse. Here, using patch-clamp electrophysiology and optogenetics in *ex vivo* mouse brain slices, we found that BLA neurons make potent synaptic connections onto MDm neurons capable of evoking action potentials. The site of this BLA input overlaps with strong innervation from locus coeruleus norepinephrine (NE) axons. We found that NE acts via α₂-adrenergic receptors to strongly reduce excitatory postsynaptic currents from BLA to MDm. NE also decreases the release probability of BLA axon terminals through a presynaptic mechanism. Postsynaptically, NE depolarizes MDm neurons and increases their tonic firing rates. These findings show that NE, whose levels are elevated during arousal and stress, can suppress transmission of affective information from BLA into MDm, thereby blunting this potent indirect pathway from BLA to mPFC.

**SIGNIFICANCE STATEMENT:** Previous anatomical studies have suggested the importance of amygdala input to the limbic thalamus. Here, using *ex vivo* electrophysiology and optogenetics in adult mice, we characterize the excitatory input from basolateral amygdala to mediodorsal thalamus, revealing the potency and physiological characteristics of this input. Further, we show that the stress-related neuromodulator, norepinephrine, binds to the α₂-adrenergic receptor to significantly dampen transmission of affective information carried by this synapse. These findings improve our understanding of key circuits involved in emotional processing and provide insight on how stress-induced neuromodulation may change circuit function, which is relevant to stress-related neuropsychiatric disorders such as depression, anxiety, schizophrenia, and PTSD.

## INTRODUCTION

The basolateral amygdala (BLA) is instrumental in the formation of emotional associations and governs cognitive and emotional processing via bidirectional communication with the medial prefrontal cortex (mPFC) (Janak and Tye, 2015; Likhtik and Paz, 2015; Mavrych et al., 2025). Dysregulation of this communication arises in neuropsychiatric disorders such as depression, anxiety, addiction, and PTSD (Wassum and Izquierdo, 2015; Ressler et al., 2022; LeDuke et al., 2023). To further understand the mechanisms underlying these processes, several studies have characterized the direct, monosynaptic connections between the BLA and mPFC (e.g. Burgos-Robles et al., 2017; Liu et al., 2020; Manoocheri and Carter, 2022). However, less is known about the indirect pathway that relays information from the BLA to the mPFC via the mediodorsal thalamus (MD), known to be part of the limbic thalamus (Vertes et al., 2015; Wolff et al., 2015). The MD and mPFC form a thalamocortical loop that underlies working memory during decision making tasks (Saalmann, 2014; Ferguson and Gao, 2018; Phillips et al., 2025) and together play key roles in behavioral flexibility (Mitchell et al., 2015; Zhang et al., 2025) and emotional stability (Rikhye et al., 2018). However, while much MD-related work has focused on its relationship with the mPFC, less is known about the inputs from BLA to MD.

Anatomical studies have established the existence of a BLA projection to MD across different organisms (McDonald, 1987; Kuroda and Price, 1991; Aggleton and Mishkin, 1984; Hintiryan et al., 2021). These studies have shown that the BLA sends an anatomically restricted input to the magnocellular sub-division of the MD (MDmc) in primates and the medial sub-division of the MD (MDm) in rats, both which are bidirectionally connected with the mPFC (Georgescu et al., 2020; Onishi et al., 2022). Recently, Ahmed and Paré (2023) found that, in rats, the majority of BLA inputs to MDm (BLA→MDm) were glutamatergic. Here, we functionally characterized this glutamatergic input from BLA→MDm. Stress- and arousal-related neuromodulators have been well established to affect activity in BLA and mPFC (Arnsten, 2009). One such neuromodulator is norepinephrine (NE), which is released by the locus coeruleus (LC) during times of stress and arousal (Poe et al., 2020). Previous studies have thoroughly characterized the role of NE in enhancing activity in BLA and mPFC, facilitating aversive emotional learning, inhibiting extinction learning, and impairing complicated cognitive processes during high-stress states (Likhtik and Johansen, 2019). NE is known to enhance the excitability of thalamocortical neurons in primary sensory thalamic nuclei (McCormick et al., 1991; Varela 2014). However, the specific effects of NE on MD and associated circuitry are unknown. Anatomically, NE-releasing fibers from LC innervate MD, and MD expresses noradrenergic α_1_ and α₂ receptors in non-human primates (Lindvall et al., 1974; Pérez-Santos et al., 2021). We hypothesized that NE may regulate the flow of information from BLA→MDm→mPFC, which may contribute to stress-induced changes in emotional and cognitive behaviors.

We examined the connection from BLA→MDm using optogenetics and patch-clamp electrophysiology in *ex vivo* brain slices from in adult mice. We find that the BLA makes potent glutamatergic synapses onto MDm neurons that are mediated by AMPA and NMDA receptors, exhibit paired pulse depression, and are capable of evoking action potentials. NE robustly inhibits these excitatory inputs via α₂-adrenergic receptor signaling, thereby reducing the vesicular release probability of BLA inputs onto MDm neurons. In addition to inhibiting BLA→MDm input, NE also enhances MDm intrinsic excitability through a non α₂-adrenergic receptor-dependent mechanism. Thus, the BLA sends strong glutamatergic inputs to MDm that likely carry important affective information. As such, stress- and arousal-related elevations in levels of NE in MDm may suppress transmission of certain types of affective information from BLA→MDm, thereby controlling amygdalo-thalamo-cortical information flow.

## MATERIALS AND METHODS

### Animals

All animal care procedures were in compliance with the NIH Guide for the Care and Use of Laboratory Animals and approved by the Institutional Animal and Care and Use Committee (IACUC) at Boston Children’s Hospital. Adult female and male C57BL/6J mice, 6 - 8 weeks of age, were ordered from Jackson Laboratory (Cat# 000664) and delivered to the animal housing facility. After stereotaxic injections, animals were moved to a smaller satellite facility. In both housing conditions, they were maintained on a 12 hour light / 12 hour dark cycle with food and water provided *ad libitum* and group-housed whenever possible.

### Stereotaxic injections

Mice were deeply anesthetized with 3% of 1 ml/s isoflurane before mounting onto a stereotaxic frame (KOPF, Model 1900) and were continuously anesthetized with 1-2.5% of 0.6 – 0.8 ml/s isoflurane throughout the procedure. The mouse’s eyes were lubricated with eye ointment and their body temperature was maintained with a heating pad. The scalp was removed of hair with Nair^TM^, sterilized with betadine and 70% EtOH, then incised to expose the skull. Craniotomies were performed with a stereotaxic drill (KOPF, Model 1911) to inject structures at the following coordinates with respect to bregma: BLA (AP = -1.6 mm, ML = ±3.3 mm, DV = -4.1 mm) or MDm (AP = -1.3, ML = + 0.3 mm, DV = -2.85 mm). The glass capillary was first lowered 0.2 mm below the listed DV coordinate, then raised to the coordinate to create a small space for injecting 150 nl of AAV2/5-Syn-ChR2-mCherry or 100 nl of Red Retrobeads (Table 1) at 1 nl/s by a nanoinjector (WPI). The micropipette was raised by 0.01 mm right after the injection and remained for 5 minutes before being slowly removed from the brain. The incision was sutured (Silk Suture, Fisher Scientific NC9140104) and treated with topical antibacterial ointment. Mice recovered on a heating pad in a clean cage before being returned to their homecage. Mice were given post-operative injections of meloxicam (1 μl/g) for 48 hours following the procedure. Mice were euthanized for slice electrophysiology experiments 4-8 weeks post-procedure or 1 week after Retrobeads injections for histology.

**Table 1.** Key resources.

| Reagent or Resource | Source | Identifier or Information |
| --- | --- | --- |
| Antibodies |  |  |
| DAPI | Invitrogen | D1306 |
| Mouse anti-NeuN | Millipore | MAB377 |
| Mouse anti-NET | Synaptic Systems | Cat# 260011 |
| Goat anti-mouse Alexa 647 | Invitrogen | a32728 |
| Streptavidin DyLight™ 488 | Invitrogen | Cat# 21832 |
| Streptavidin Alexa Fluor™ 647 | Invitrogen | Cat# S32357 |
| Chemicals |  |  |
| Neurobiotin® Tracer | Vector Laboratories | SP-1120-50 |
| Alexa Fluor™ 488 Hydrazide | Thermo Fisher Scientific | A10436 |
| Alexa Fluor™ 647 Hydrazide | Thermo Fisher Scientific | A20502 |
| Methoxyverapamil | Sigma-Aldrich | M5644 |
| Tetrodotoxin | Tocris | Cat# 1078 |
| 4-Aminopyridine | Tocris | Cat# 0940 |
| Bicuculline | Tocris | Cat# 0130 |
| NBQX | Tocris | Cat# 0373 |
| CPP | Tocris | Cat# 0247 |
| Noradrenaline bitartrate | Tocris | Cat# 5169 |
| Yohimbine hydrochloride | Tocris | Cat# 1127 |
| UK14304 tartrate | Tocris | Cat# 2466 |
| Stereotaxically Injected Reagents |  |  |
| AAV2/5-Syn-ChR2-mCherry | Boston Children's Hospital<br>Viral Core | 3.86 x 10<sup>13</sup> vg/mL |
| Red Retrobeads™ | Lumafluor Inc. |  |

### Slice preparation

Mice were euthanized for patch clamp electrophysiology experiments during the middle of the light cycle. Mice were deeply anesthetized with isofluorane and transcardially perfused with a chilled, oxygenated K-Gluconate based solution comprised of the following (in mM): K-Gluconate 130, KCl 15, EGTA 0.05, HEPES 20, glucose 25, and adjusted to pH of 7.3 - 7.4 (with KOH) and an osmolarity of 310-315 mOsm. The brain was dissected and removed from the mouse, and the cerebellum was trimmed off with a steel razor blade. Parts of the left and right hemispheres containing the BLA, ∼2-3 mm in width, were trimmed off sagittally, using the lateral edges of the olfactory bulb as a visual aid. Further processing of tissue containing the BLA is further described in the section *Tissue preparation and immunohistochemistry*. The rest of the brain containing MD was sagittally sectioned into 250 um slices using a sapphire blade (Delaware Diamond Knives) on a vibratome (VT1200S; Leica), using the previously described K-Gluconate based solution as an oxygenated, chilled cutting solution. Slices containing MD were collected and deposited into oxygenated artificial cerebrospinal fluid (ACSF) at 33°C to recover for 15 minutes. The ACSF comprised of the following (in mM): 125 NaCl, 2.5 KCl, 1.25 NaH2PO4, 26 NaHCO3, 1 MgCl_2_, 2 CaCl_2_, 25 mM glucose, adjusted to pH of 7.3 – 7.4 (with HCl) and 310 - 315 mOsm. After the 15-minute period, the slices were transferred to room temperature (RT) and allowed a further acclimation period of 45 minutes. Reagents were purchased from Sigma-Aldrich.

Specifically, with reference to the sagittal Allen Brain Atlas and the Paxinos Atlas, the habenulo-interpeduncular tract (fr) innervating the lateral habenula was used as a visual guide to collect slices containing the MD, but not the paraventricular thalamus (PVT), at ± 0.2 – 0.35 mm lateral from the midline (Figure S3B). The emergence of the medial terminal nucleus of the accessory optic tract (mt) innervating the anteromedial nucleus of the thalamus (AM) was used to signify more lateral slices (± 0.35 - 0.48 mm from the midline) containing MD (Figure S3C). Slices containing the PVT (± 0 – 0.15 mm from the midline) were collected but were not used for recording experiments (Figure S3A).

### Patch clamp electrophysiology and optogenetic stimulation

Whole-cell recording pipettes were pulled with a horizontal electrode puller (P-97, Sutter Instruments) from glass capillary tubes (Drummond Scientific Company, 2-000-100). For voltage clamp experiments, 2 - 4 MΩ resistance pipettes were used and filled with a CsCl based internal recording solution consisting of (in mM): 35 CsF, 100 CsCl, 10 EGTA, 10 HEPES, and the L-type calcium channel antagonist, 0.1 methoxyverapamil (Table 1), adjusted to pH 7.3 (with CsOH) and 290 mOsm. For current clamp experiments, 4 - 5 MΩ pipettes were used and filled with a K-gluconate based internal recording solution comprised of (in mM): 135 K-gluconate, 7 KCl, 0.5 EGTA, 10 HEPES, 10 mM Na_2_-phosphocreatine, 4 mM Mg_2_-ATP, 0.4 mM Na-GTP. In most experiments, the internal included 0.2% Neurobiotin^®^ Tracer and/or 20 μM Alexa Fluor^TM^ 488 or 647 Hydrazide (Table 1) to visualize patched neurons. Reagents were purchased from Sigma-Aldrich.

Freshly oxygenated ACSF was perfused onto the slice at RT with a flow rate of 1.6 - 2 ml / min. For all voltage clamp experiments, 20 μM bicuculline was used to block GABA_A_ mediated inhibitory transmission. For experiments to isolate monosynaptic currents, 0.5 μM tetrodotoxin and 1 mM 4-aminopyridine were used. For current clamp experiments to measure intrinsic excitability, 20 μM NBQX, 20 μM CPP, and 20 μM bicuculline were used to block AMPA, NMDA, and GABA_A_ receptors, respectively. For noradrenergic pharmacology experiments, the following were used: 10 μM noradrenaline bitartrate, 20 μM yohimbine hydrochloride, and 20 μM UK14304 tartrate (Table 1).

Full-field optogenetic stimulation of ChR2-expressing BLA axon terminals was performed to obtain the maximal optogenetically evoked excitatory post synaptic currents (oEPSCs). A CoolLED pE unit supplied 470 nm light through a 60x objective (Olympus LUMplanFL N 60x/1.00W), and brief 0.2 ms pulses of light were used with a maximal intensity of 10 mW/mm^2^. Trials holding at -70 and +40 mV were alternated with an inter-trial interval of 30 seconds for voltage clamp experiments. For paired-pulse experiments, inter-stimulus intervals of 250, 500, 750, and 1000 ms were randomly sampled, and paired-pulse ratio was calculated as the amplitude of the second EPSC divided by that of the first. For current clamp experiments related to Figure 7, a constant amount of holding current was applied to maintain a membrane potential of -67 ± 3 mV, and 1 second current steps were applied from -75 to 150 pA in 25 pA steps to assess burst and tonic firing. For experiments related to Figures 3, the same K-gluconate based internal recording solution was used with 4 - 5 MΩ pipettes. Recording paradigms began first in voltage clamp, then were switched to current clamp at the resting membrane potential (RMP) of the neuron which ranged between -70 mV to -57 mV and corresponded to the tonic mode (Figure 3Aii and 3C). The minimal light stimulation required to elicit BLA-evoked spikes in MDm neurons was determined at RMP in the tonic mode. The light stimulation intensity was reduced from a maximal power of 10 mW/mm^2^ to minimal power in -50% increments of the previously used intensity. If a reduced light power was not sufficient to induce a spike, light power was raised to the halfway point between the previous successful step and the unsuccessful step. Once the minimal light power was determined in current clamp, the recording configuration was switched to voltage clamp and the amplitude of BLA→MDm oEPSCs evoked by this light power were determined. After optogenetically stimulating BLA input at these potentials, most MDm neurons naturally hyperpolarized to burst mode, which ranged from -77 mV to -73 mV.

Data were acquired using an Axopatch 200B amplifier, a BNC-2090A National Instruments terminal block, and Igor Pro 8 software. Recordings were collected at 20 kHz and low-pass filtered at 5 kHz (current clamp) or 1 kHz (voltage clamp). Series resistance and input resistance were calculated using seal tests of a -5 mV step (voltage clamp) or -25 pA injection (current clamp) before each trial of the recording protocol. For current clamp data, the liquid junction potential was experimentally determined (-12.3 mV) and compensated for post-hoc. Neurons that exhibited series resistance greater than 10 MΩ and changed by more than 10% across voltage clamp experiments were excluded unless otherwise indicated.

### Single fiber stimulation

When preparing sagittal slices containing MD for single fiber stimulation experiments, we found a parasagittal cut made with a 5-15° angle with respect to the midline on the hemisphere opposite from that being recorded enhanced our ability to record single BLA fibers to MDm. The parasagittal cut helped to preserve more of the ventral amygdalofugal tract within the slice, as the reconstructed amygdala to MDm input showed that the amygdala projection traveled in a very slight medial to lateral direction from the anterior to posterior axis in MDm (Figure 1D, top). Beyond this parasagittal cut, slices were prepared as described above and only 1 hemisphere was used per animal. The hemisphere containing the relevant BLA injected with AAV2/5-Syn-ChR2-mCherry was trimmed off with a steel blade and postfixed for post-hoc histology as described above.

**Figure 1.**
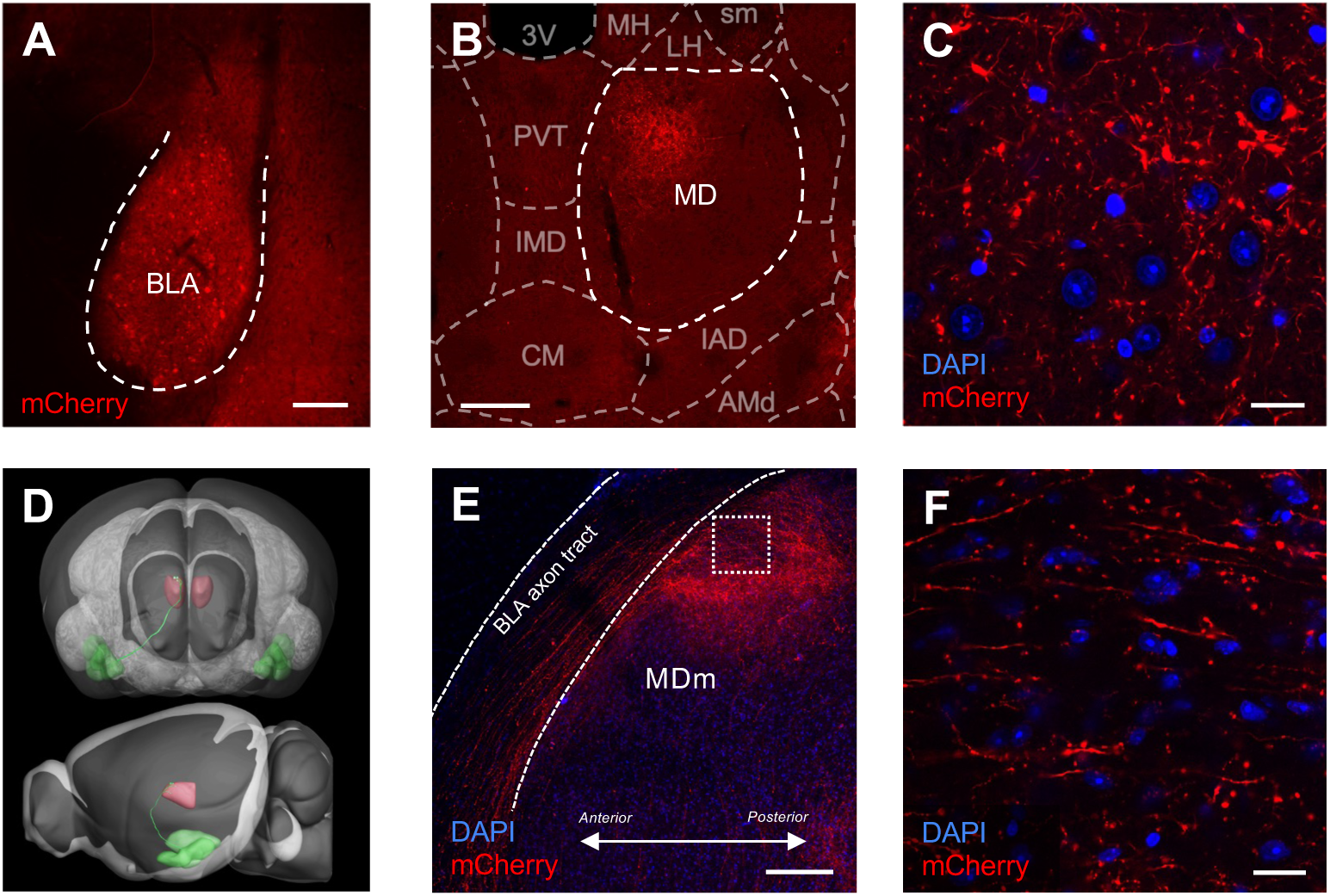
BLA sends inputs to MDm. ***A***, Image showing a coronal section of the BLA following injection of AAV2/5-Syn-ChR2-mCherry. Cell bodies labeled with mCherry were observed (red). Scale bar = 200 μm. ***B***, BLA axons and axon terminals were observed in a coronal section of medial MD (MDm). Allen Brain Reference Atlas overlaid to show anatomical structures. Scale bar = 200 μm. ***C***, BLA axon terminals (mCherry, red) in MDm (DAPI, blue) from ***B*** at higher magnification. Scale bar = 20 μm. ***D***, Reconstruction of a virally labeled axonal projection from amygdala (green) to mediodorsal thalamus (MD, red) from the Allen Brain Mouse Connectivity dataset, Experiment 125832322. Top, coronal plane; bottom, sagittal plane. ***E***, BLA was injected with AAV2/5-Syn-ChR2-mCherry as in Figure 1A. A sagittal slice of MDm preserved a portion of the BLA axon tract (mCherry, red) that innervates MDm (DAPI, blue). Scale bar = 200 μm. ***F***, Higher magnification of MDm area outlined by the dashed square in ***E***. BLA axons (mCherry, red) travelling across MDm from anterior (left) to posterior (right). Scale bar = 20 μm. AMd, anteromedial nucleus, dorsal part; BLA, basolateral amygdala; CM, central medial nucleus of the thalamus; IAD, interanterodorsal nucleus of the thalamus; IMD, intermediodorsal nucleus of the thalamus; LH, lateral habenula; MH, medial habenula; PVT, paraventricular thalamus; sm, stria medullaris; 3V, third ventricle.

To optogenetically stimulate single ChR2-expressing BLA axons within the ventral amygdalofugal tract in our sagittal slice preparation, we used a 200 μm-thick optic fiber, 0.39 nA (ThorLabs), threaded through a glass pipette. The fiber was positioned 200+ μm away from the ventral amygdalofugal tract for single fiber experiments as illustrated in Figure 2H. 470 nm light stimulation was provided by a fiber-coupled LED (ThorLabs, M470F3). Single fibers were identified by modifying the failures method previously described (see supplemental materials and Figure S2 of Noutel et al., 2011). Briefly, opto-fiber trials were similar to those used for previously described voltage clamp experiments, using brief pulses of 0.2 ms 470 nm light with a 250 ms ISI at -70 mV HP to record primarily AMPAR-mediated currents. The optogenetic stimulation intensity was incrementally reduced to achieve a 20 - 50% success rate of evoking a fiber-evoked excitatory postsynaptic current (fEPSC) on the first pulse of light. Gradual reduction in stimulation intensity entailed initially reducing light power from 10 mW/mm^2^ to 0.1 mW/mm^2^ in 8 -50% increments. 100% success rate, sometimes with varying fEPSC amplitudes, was still observed even with minimal light power with 0.2 ms light stimulations. Thus, the optogenetic stimulation duration was further reduced from 0.2 to 0.0175 - 0.018 ms. For example, in Figure 2I, there were some failures to evoke an fEPSC, some successes only on the second pulse, and some successes for both pulses of optogenetic stimulation. The amplitude of fEPSCs for successful first pulses did not vary significantly between trials. These parameters support the idea that these recordings are from a single fiber.

**Figure 2.**
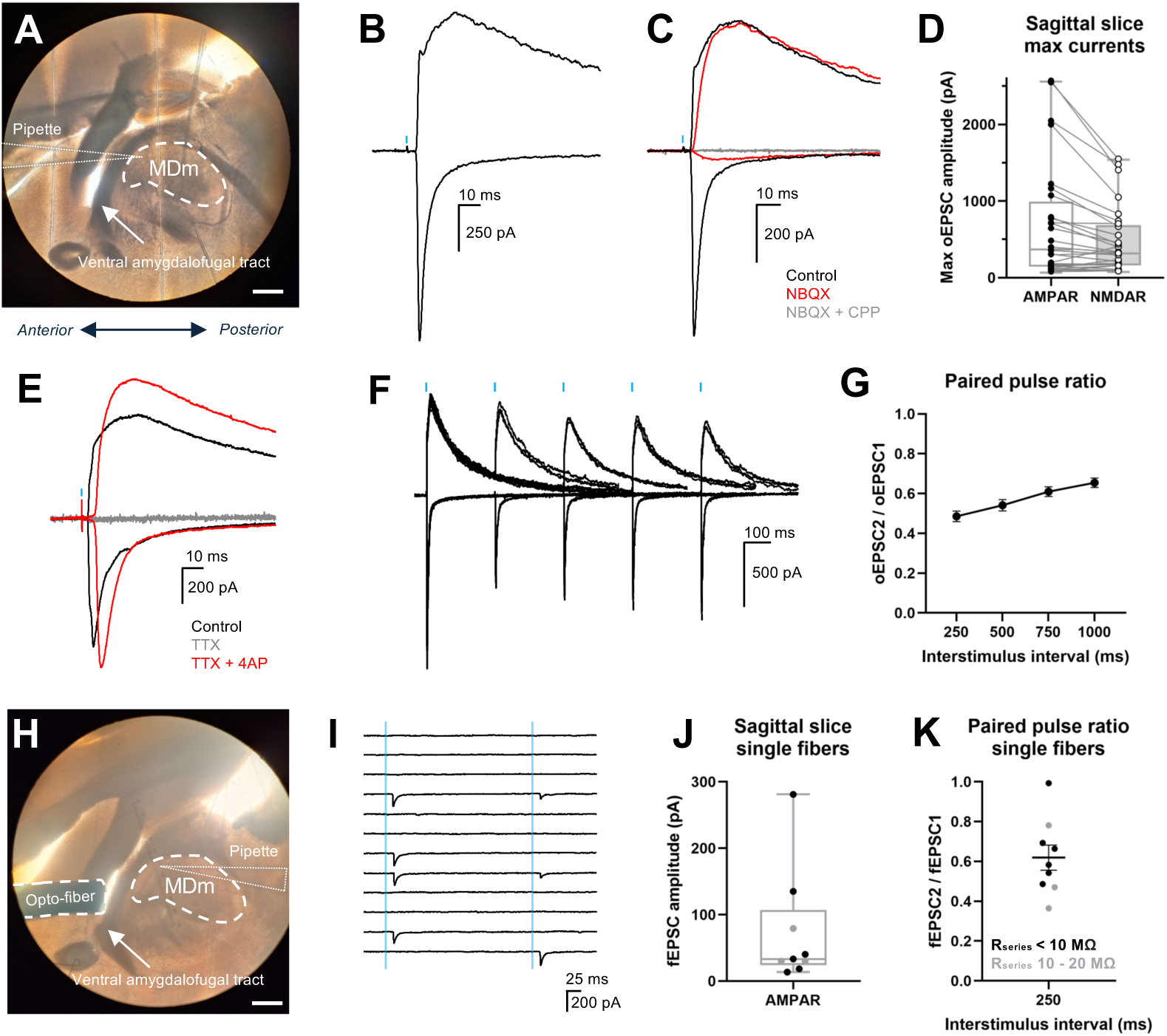
BLA sends strong excitatory input to MDm. ***A***, Sagittal slice preparation of MDm shown under DIC for patch clamp electrophysiology, preserving the ventral amygdalofugal tract that carries inputs from BLA→MDm (arrow). Scale bar = 500 μm. ***B***, Optogenetically-evoked excitatory postsynaptic currents (oEPSCs) from BLA→MDm inputs by 0.2 ms 470 nm light stimulation (blue bar) following AAV2/5-Syn-ChR2-mCherry injection into BLA. Example oEPSCs are shown at two holding potentials (HP):-70 mV (bottom, largely AMPAR-mediated current) and +40 mV (top, AMPAR- and NMDAR-mediated current). ***C***, oEPSCs recorded at HP = -70 and +40 mV after application of 20 μM NBQX to block AMPA receptors (red), and after additional application of 20 μM NBQX + 50 μM CPP to block AMPA and NMDA receptors (grey). N = 3 cells/3 mice. ***D***, Plot of BLA→MDm peak AMPAR and NMDAR oEPSC amplitudes evoked by maximal light stimulation (10 mW/mm^2^, 0.2 ms duration). oEPSC responses from individual neurons are connected by grey lines. Amplitudes ranged from 65 pA to 2.6 nA with a median of 366.85 pA and 318.45 pA for AMPAR and NMDAR, respectively. Box, 25% - 75% IQR. N = 28 cells/19 mice. ***E***, Application of 0.5 μM tetrodotoxin (TTX) abolishes oEPSCs recorded at HP =-70 and +40 mV (gray). oEPSCs are recovered with bath application of 0.5 μM TTX with 1 mM 4-aminopyridine (4AP) (red), demonstrating that these oEPSCs are monosynaptic. N = 3 cells/3 mice. ***F***, Superimposed pairs of BLA→MDm oEPSCs recorded at HP = -70 and +40 mV with varying interstimulus intervals (ISI); 250, 500, 750, and 1000 ms from one MDm neuron (all paired pulse trials overlaid, 3 trials/ISI per HP). ***G***, Paired pulse ratio (PPR) of each neuron in the dataset was calculated as the ratio of oEPSC2/oEPSC1 peak amplitudes at different ISIs. Average PPR ± SEM is shown. N = 15 cells/12 mice. ***H,*** Sagittal slice preparation of MDm as in ***A.*** An optic fiber (outlined) was positioned close to the ventral amygdalofugal tract (arrow) to generate an optic fiber-evoked EPSC, or “fEPSC.” Scale bar = 500 μm. ***I***, Example fEPSC responses to consecutive trials of stimulation of a BLA→MDm input obtained from the slice shown in ***H.*** Each trial shows the response to pairs of minimal optogenetic stimulation (blue bar, 250 ms ISI, inter-trial interval of 30 sec). Minimal stimulation was defined using the failures method (Noutel et al., 2011; see methods). ***J***, Amplitudes of single BLA fibers into MDm, as obtained with paradigm shown in ***I***. Median AMPAR peak fEPSC amplitude was 33.41 pA. Box, 25% - 75% IQR. ***K,*** Average PPR ± SEM of single fibers from ***J*** with a 250 ms ISI. ***B, C, E, F, I***: Blue bars indicate time point of optogenetic stimulation. ***J, K***: N = 9 cells/5 mice. Only recordings from neurons with <10 MΩ series resistance were analyzed unless otherwise noted.

**Figure 3.**
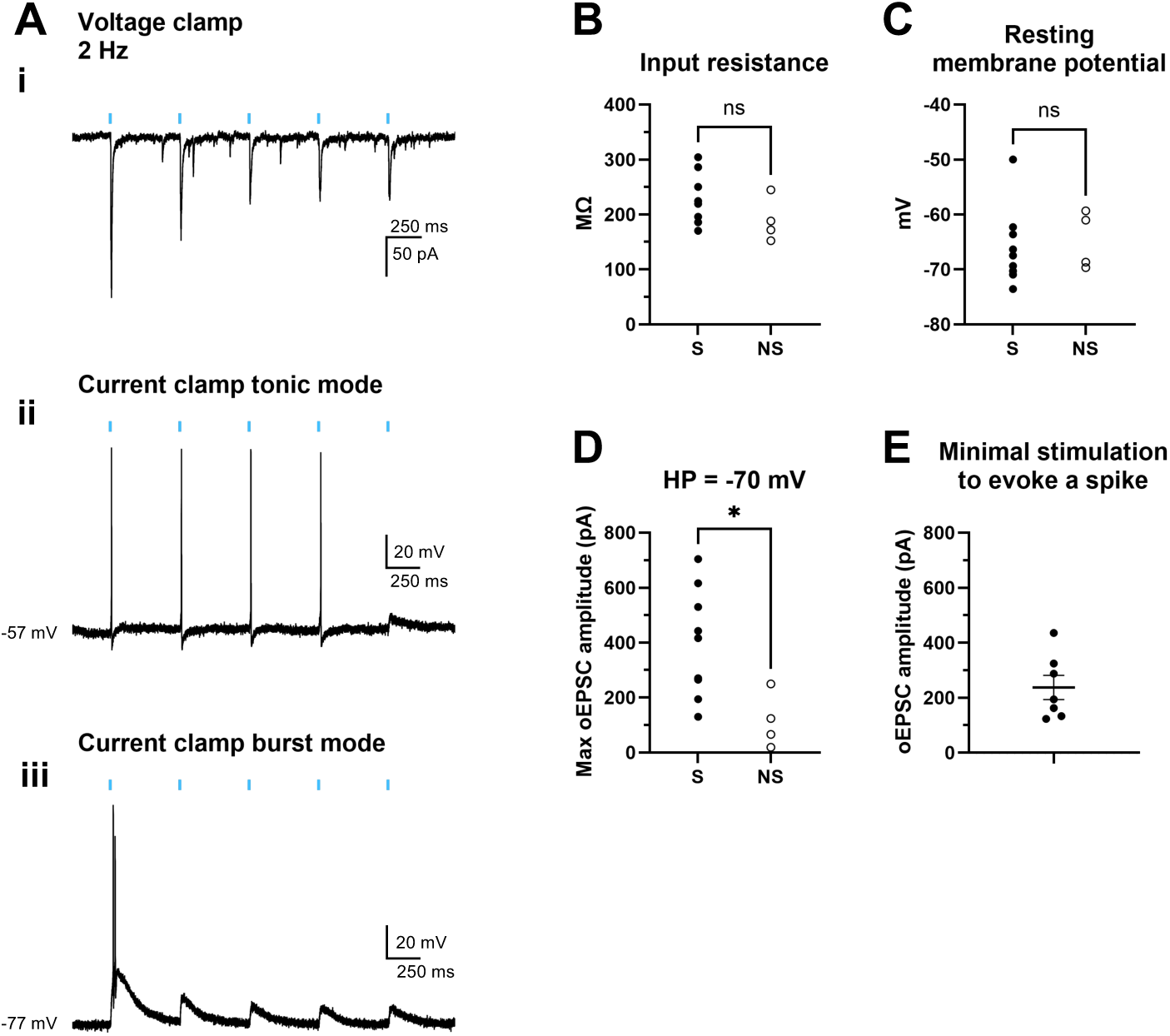
Optogenetic stimulation of BLA input evokes spiking in a subset of MDm neurons. *Ex vivo* recordings from sagittal slices of MDm in mice expressing with AAV2/5-Syn-ChR2-mCherry in BLA. No synaptic blockers were used. ***A***, BLA inputs were stimulated with 5 0.2-ms pulses of 10 mW/mm^2^ 470 nm light (blue bars) at 2 Hz while recording in voltage clamp (*i*), current clamp (tonic mode) at resting membrane potential (RMP) (*ii*), and current clamp (burst mode) at hyperpolarized potentials between -77 and -75 mV (*iii*) in the same MDm neuron. ***B-C***, Input resistances and RMPs of MDm neurons in which BLA inputs either evoked spiking (S) or did not evoke spiking (NS)*. **B***, Input resistances between the S and NS groups were not significantly different (unpaired t-test, t_(11)_ = 1.366, p = 0.1994). ***C,*** RMPs between the S and NS groups were not significantly different (unpaired t-test, t_(11)_ = 0.3342, p = 0.7445). ***D***, Max oEPSC amplitudes of BLA inputs obtained at HP = -70 mV that either evoked spiking (S) or did not evoke spiking (NS) at RMP in MDm neurons. oEPSC amplitudes were significantly lower for the NS group compared to the S group (unpaired t-test, t_(11)_ = 2.675, p = 0.0216). ***E***, For MDm neurons from (S) group in ***D***, light power was titrated down to the minimal stimulation sufficient to evoke a spike. The peak amplitudes of BLA→MDm oEPSCs obtained at this minimal stimulation level are plotted for each neuron (mean ± SEM = 237 pA ± 43.86 pA; HP = -70 mV). ***B-D***: N = 12 cells / 4 mice.

### Tissue preparation and immunohistochemistry

Mice used for Figures 1 and S3 were euthanized with an intraperitoneal injection of pentobarbital. Their brain tissue was fixed by transcardial perfusion and overnight fixation with 4% paraformaldehyde (Electron Microscopy Sciences) and rinsed with 1xPBS before sectioning into 60 - 100 μm thick sections on a vibratome (Leica VT1000). For Figure S2, sections were incubated in blocking solution of 5% goat serum with 0.1% Triton in 1xPBS (PBS-T) for 1 hour at RT, then incubated for 1-2 days at 4°C in primary antibody solution containing either mouse anti-NeuN (1:1000) or mouse anti-NET (1:1000) in 2% goat serum and 0.1% PBS-T (Table 1). Sections were then washed with 0.1% PBS-T and incubated for 1-2 hours at RT in a secondary antibody solution containing Alexa 647 goat anti-mouse antibody (1:1000) and washed with 1xPBS (Table 1). For both Figure 1 and S1, sections were stained for 5-10 minutes in DAPI (1:1000) in PBS, rinsed with 1xPBS, and mounted onto slides (VWR Superfront Plus, 48311-703) with Vectashield^®^ Antifade Mounting Medium (Vector Laboratories, H-1000).

For processing slices that underwent patch-clamp electrophysiology such as those from Figure 4A and C, slices were drop-fixed in 4% paraformaldehyde, incubated overnight, then rinsed with 1xPBS. To prepare for immunostaining, slices were cryoprotected in 30% sucrose in 1xPBS for at least 2 hours at 4°C. The sucrose solution was removed and slices were quickly frozen on dry ice and thawed for freeze-thaw permeabilization. Slices were blocked in a solution of 5% goat serum in 0.3% PBS-T for 1 hour at RT. They were then incubated for 2-3 days in a primary antibody solution of mouse anti-NET (1:1000) in 2% goat serum with 0.3% PBS-T, then rinsed with 0.3% PBS-T and incubated for 1 hour at RT in a secondary antibody solution containing Alexa 647 goat anti-mouse antibody (1:1000) and Streptavidin DyLight^TM^ 488 (1:1000). Subsequently the slices were washed with 0.3% PBS-T, immunostained against DAPI (1:1000 in 1xPBS) for 5-10 minutes, then rinsed with 1xPBS. Slices were mounted on glass slips as previously described. Slices were imaged with confocal microscopy (Zeiss LSM 700, Leica TCS SP8 for tilescans) and referenced with both the sagittal Allen Brain Atlas and the Paxinos Atlas to confirm that labeled neurons were in MD.

**Figure 4.**
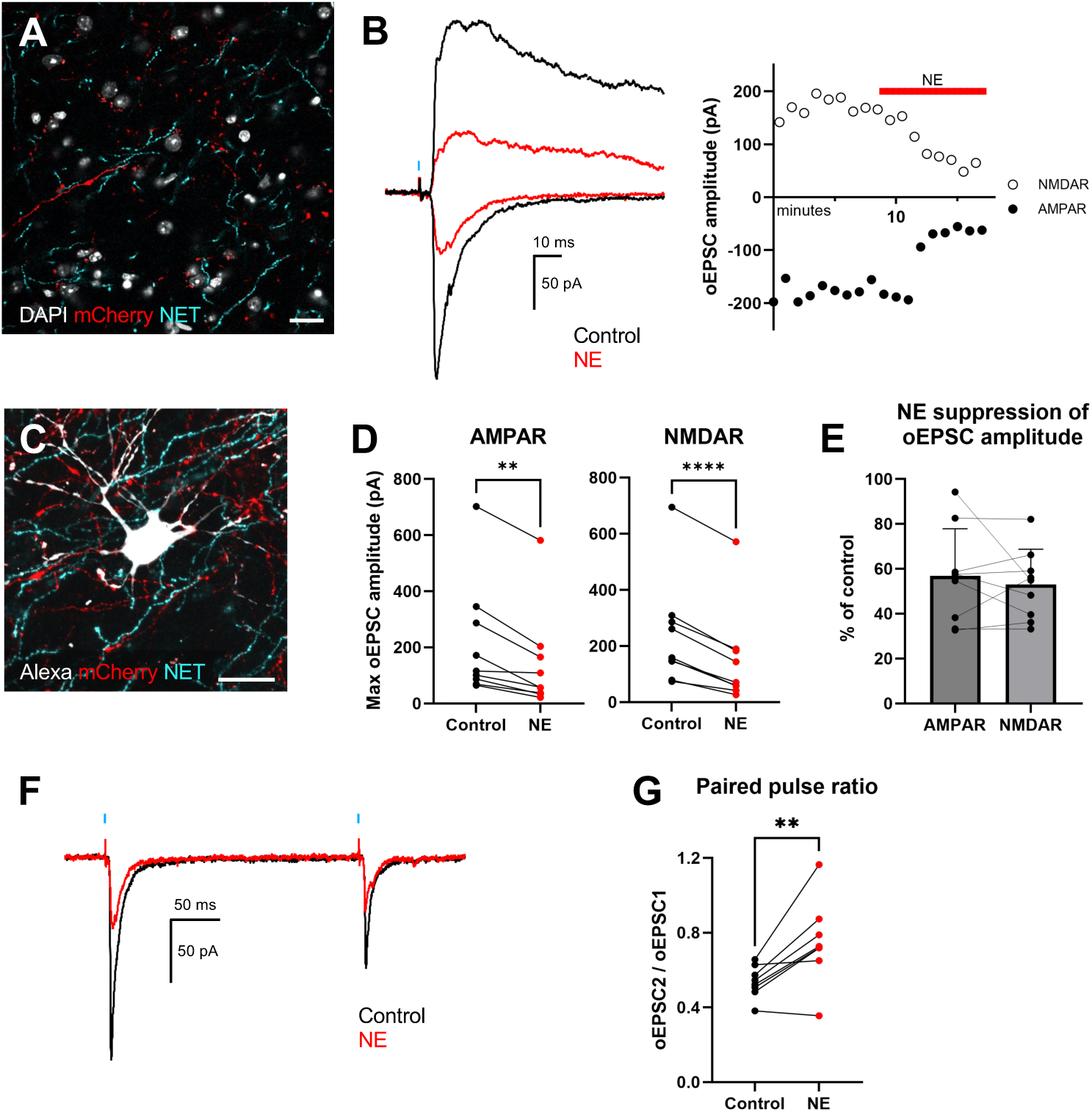
Norepinephrine (NE) reduces the amplitude and increases the paired pulse ratio of BLA→ MDm oEPSCs. ***A***, BLA was injected with AAV2/5-Syn-ChR2-mCherry to label BLA axons (red). Immunostaining for the norepinephrine transporter (NET) was performed to label NE-releasing axons (cyan) in a sagittal section of MDm. Scale bar = 20 μm. ***B***, Example oEPSCs from BLA→MDm inputs, shown at HP = -70 mV (inward current, largely AMPAR-mediated current) and HP = +40 mV (outward current, AMPAR- and NMDAR-mediated current). 10 μM NE reduced the amplitudes of both AMPAR and NMDAR currents. Black = control, red = NE application. Blue bar indicates time point of 0.2 ms light stimulation. Right, time-course of oEPSC peak amplitude. Black circles = AMPAR, white circles = NMDAR, red bar = duration of 10 μM NE application. ***C***, Example of a patched cell filled with Alexa Fluor 488 dye (white), which exhibited inhibition of BLA to MDm peak oEPSC amplitudes in response to bath application of 10 μM NE. BLA inputs (labeled by viral mediated expression of mCherry in BLA) intermingle with NET positive inputs proximal to the filled cell. Scale bar = 20 μm. ***D***, NE significantly reduced the peak amplitudes of each neuron’s oEPSCs for AMPAR-(t_(8)_=4.561, p=0.0018) and NMDAR-(t_(8)_=8.420, p=<0.0001) mediated currents. Lines connect values from the same cell. ***E***, oEPSC amplitudes from ***D*** during NE application normalized as a % of pre-drug control. NE application reduced AMPAR and NMDAR oEPSCs to 57.18% ± 21.85% and 54.04% ± 16.04% of their original pre-drug amplitude. Lines connect values from the same cell. ***F***, Example AMPAR oEPSCs from BLA→MDm inputs in response to pairs of stimuli with a 250 ms ISI. 10 μM NE reduced the amplitudes and increased the paired-pulse ratio of the displayed oEPSCs compared to pre-NE administration. Black = control, red = NE application. ***G***, NE significantly increased the paired pulse ratio of BLA→MDm oEPSCs from 0.52 ± 0.10 to0.72 ± 0.23, t_(8)_ = 4.034, p=0.0038. Lines connect values from the same cell. *Statistics (**D, G**)*: Pairwise t-tests. (***D, E, G***): Values are mean ± SD. N = 9 cells/9 mice.

### Posthoc validation of stereotaxic injection sites

To validate AAV2/5-Syn-ChR2-mCherry infection in BLA (and not other areas such as the ventral hippocampus), ∼2-3 mm lateral sections of the left and right hemispheres containing the BLA were obtained as described above in *Slice preparation.* These lateral sections were drop-fixed and incubated overnight in 4% paraformaldehyde, then rinsed 3x with 1xPBS the next day. To prepare for sectioning, tissue was embedded in 3% agarose gel. The embedded tissue was then oriented coronally, and 100 μm thick coronal sections were obtained using a vibratome (Leica VT1000), which were then mounted onto slides (VWR Superfront Plus, 48311-703) with VECTASHIELD^® PLUS^ Antifade Mounting Medium with DAPI (Vector Laboratories, H-2000). Tilescan images of these slices were obtained using an epifluorescent microscope (Zeiss Axio Imager.Z2) to visualize mCherry and DAPI. The BLA was identified as being within the amygdalar capsule, which was marked by an increased density in DAPI signal (Figure S1). Data from hemispheres in which the BLA had not been infected with mCherry, or in which mCherry signal extended significantly beyond the amygdalar capsule, were excluded from the dataset.

### Experimental design and statistical analyses

Experiments were performed with alternating cohorts of male and female mice to ensure equal sampling from either biological sex, with no significant differences found in data between either sex. 1-2 slices were used from each mouse and were sampled from either brain hemisphere, as the BLA to MDm projection was confirmed to be ipsilateral. Once slices were exposed to drugs, they were not re-used to prevent drug re-exposure contaminating experimental results. Therefore, we obtained one neuron from each slice for experiments requiring drug applications.

Statistical tests were performed using GraphPad Prism 10.5.0. For datasets that passed the Kolmogorov-Smirnov (KS) normality test, mean ± SEM values are shown. For datasets that did not pass the KS normality test, median values were determined instead. *P* values < 0.05 were considered statistically significant. For pharmacological electrophysiology experiments, two-tailed t-tests were used to compare two groups with different drug conditions and were paired if data were compared from the same neuron pre and during drug application. For experiments with more than 2 groups such as firing rate curves, a repeated-measures ANOVA was used to determine statistically significant differences due to drug applications. If there were significant differences, post-hoc Tukey tests were used to determine which groups were statistically significantly different from each other. A mixed ANOVA was used instead of a repeated measures ANOVA if groups did not consist of the same N, as indicated in the figure legends.

## RESULTS

### A novel brain slice preparation for examining BLA**→**MDm inputs

We first sought to label the BLA→MDm projection in adult mice. Injections of AAV2/5-Syn-ChR2-mCherry in the BLA resulted in robust labeling of mCherry+ BLA axons terminating in the medial sub-region of the mediodorsal thalamus (MDm) (Figure 1A-B), often close to DAPI-stained MDm cell bodies (Figure 1C, Figure S1A,B). To confirm that BLA (and not merely peri-BLA regions) sends inputs to MDm, we injected red retrobeads into MDm (Figure S2A). The retrobeads retrogradely labeled various neurons within the BLA, including the region that had initially been labeled with ChR2 in Figure 1A (Figure S2B-D). The habenula, which was also labeled with retrobeads above MDm, does not receive inputs from the BLA (Hintiryan et al., 2021). These results confirmed previous anatomical studies describing the BLA→MDm projection in the rodent brain (McDonald, 1987; Hintiryan et al., 2021).

We next developed an acute brain slice preparation to record optogenetically evoked postsynaptic currents from BLA→MDm. To this end, we were interested in creating a slice preparation preserving the trajectory of labeled BLA projections to facilitate identification of innervated MDm neurons. To determine the optimal orientation for this brain slice, we reconstructed the amygdala→MD projection from data on the Allen Brain Atlas’s Mouse Connectivity repository (Experiment 125832322) that labeled amygdala axonal projections with an injection of AAV-EGFP into the basolateral and basomedial amygdala. These reconstructions revealed that the amygdala sends substantial inputs to MDm via a distinct axonal bundle, which comprises part of the ventral amygdalofugal pathway and is one of the primary outputs of the BLA (Figure 1D). Notably, as the amygdalar axon tract approaches MDm, it projects parallel to the sagittal plane. Thus, we cut brains sagittally from mice that had been infected in the BLA with AAV2/5-Syn-ChR2-mCherry. This sagittal angle successfully preserved part of the distal end of the amygdala axon bundle containing mCherry+ BLA axons (Figure 1E), similar to how parasagittally cut brain slices containing the dorsal lateral geniculate nucleus (dLGN) can preserve the optic tract (Chen and Regehr, 2000; Litvina and Chen, 2017). Additionally, the sagittal angle captures the defasciculation of the BLA axons from the ventral amygdalofugal pathway as they project from the anterior to the posterior direction and arborize in the MDm (Figure 1E-F, Figure S3).

A prior study showed that the majority of BLA→MDm inputs were glutamatergic in the rat (Ahmed and Paré, 2023). We further examined the synaptic properties of these inputs by performing whole-cell voltage clamp recordings of MDm neurons while optogenetically stimulating the BLA inputs (full-field 470 nm light stimulation, 0.2 ms duration, through a 60x microscope objective). Recordings were performed in the presence of 20 μM bicuculline in the external recording solution to block inhibitory transmission and used two holding potentials: (a) - 70 mV to record primarily AMPAR-mediated currents, and (b) +40 mV for AMPAR- and NMDAR-mediated currents. Resulting optogenetically-evoked excitatory postsynaptic currents (oEPSCs) varied in amplitude across both holding potentials, ranging from 65 pA to 2.6 nA (Figure 2B, D). To confirm that these currents were mediated by AMPA and NMDA receptors, we applied 20 μM NBQX, an AMPA receptor antagonist, which significantly abolished the fast transient component of the oEPSC obtained at both -70 mV and +40 mV. Additional application of 50 μM CPP, an NMDA receptor antagonist, together with NBQX completely abolished both oEPSCs at -70 mV and +40 mV (Figure 2C). In separate experiments, oEPSCs were also abolished with bath application of tetrodotoxin (TTX, 0.5 μM), supporting the idea that these evoked currents were action potential dependent. Addition of 4-aminopyridine (4AP, 1 mM) together with TTX resulted in the recovery of the oEPSC, consistent with a monosynaptic input from BLA→MDm (Figure 2E) (Yamawaki et al., 2016).

We also examined the BLA→MDm synaptic response to pairs of stimulation pulses. We assessed the paired-pulse ratio (PPR = peak amplitude of oEPSC2/oEPSC1; HP: -70 mV) at 4 different interstimulus intervals (ISIs, Figure 2F). Robust paired pulse depression was observed even with ISIs of 1000 ms (Figure 2G). We asked whether the observed paired-pulse depression to full-field optogenetic stimulation could be an artifact of ChR2 desensitization, leading to dropout of axon fibers evoked by the second light pulse. To rule out this possibility, we used an opto-fiber to stimulate the amygdala axon bundle far from the patched neuron of interest to induce fiber-evoked excitatory postsynaptic currents (fEPSCs) and examined the paired-pulse response to activation of isolated single BLA fibers onto an MDm neuron. We titrated the light power to isolate a single fiber using the “failures” method (Figure 2I; see also methods from Noutel et al., 2011; Litvina and Chen, 2017). Isolated individual BLA fibers also exhibit paired-pulse depression to a 250 ms ISI optogenetic stimulation (Figure 2I, K). Amplitudes of single BLA fibers ranged from 10 - 300 pA (Figure 2J). This EPSC magnitude was often just a fraction of the maximal axon-evoked EPSC, suggesting that in some instances, multiple BLA inputs converge on a single MDm neuron to contribute to the large net excitatory drive.

### BLA inputs can drive action potential firing in MDm neurons

Thalamocortical neurons have two modes of responding to stimuli depending on their membrane potential. The first mode is the tonic mode, which occurs at more depolarized potentials and consists of singular action potentials fired directly in response to an excitatory input. The second mode is the burst mode, occurring at more hyperpolarized potentials, in which neurons fire a burst of spikes in response to depolarization (Sherman, 2001). We were curious how these BLA inputs contribute towards postsynaptic potentials or spiking activity in MDm neurons in the tonic and burst modes. To this end, we compared the synaptic responses to the action potential (AP) responses to optogenetic stimulation of BLA inputs. After recording the oEPSCs in voltage clamp (Figure 3Ai), we then switched into current clamp and recorded changes in membrane potentials in tonic mode (Figure 3Aii) and burst mode (Figure 3Aiii) in response to the same light stimulation. These recordings were performed without bicuculline or other synaptic blockers, as we wanted to observe the overall effect of BLA→MDm inputs with inhibitory transmission intact in our preparation. We found that optogenetic stimulation of BLA input was sufficient to evoke AP firing in 8/12 MDm neurons at resting membrane potential (RMP). We compared the properties of the neurons with (S) and without (NS) spiking and did not find any differences in input resistance or resting membrane potential (Figure 3B, C). However, the peak amplitudes of BLA→MDm oEPSCs that were able to evoke a spike were significantly greater when compared to those that were unable to evoke a spike (Figure 3D). For neurons in the (S) group, we titrated the light power to find the minimum stimulation necessary to evoke a spike at RMP in current clamp (see Methods for more details). We then switched to voltage clamp to record the peak amplitude of the BLA→MDm oEPSC evoked by this minimal stimulation at HP = -70 mV. On average, a peak oEPSC of 237 ± 43.86 pA could drive AP firing in MDm neurons (Figure 3E).

We next asked how synaptic depression to trains of stimulation affect AP firing at BLA→MDm connections. We found that the likelihood of spiking was highest in response to the first stimulation of a train and then decreased with subsequent stimulations. Figure S4 compares spiking to higher frequency stimulation (5 and 10 Hz). The probability of spiking decreases with increasing stimulation frequency in the tonic mode (Figure S4B), while the number of BLA-evoked bursts remained consistent across 2, 5, and 10 Hz in the burst mode (Figure S4C). These results suggest that summation of sub-threshold EPSPs does not contribute significantly to spiking at these frequencies.

Altogether, our results confirm that the BLA makes excitatory glutamatergic synapses onto MDm, many of which are strong enough to evoke spiking in the tonic and burst modes. Furthermore, BLA→MDm inputs exhibit robust paired-pulse depression, suggesting that they have a high probability of neurotransmitter release.

### Noradrenergic modulation of the BLA**→**MDm synapse

We found that noradrenergic neurons in the locus coeruleus (LC) were robustly labeled following injection of retrobeads in MDm, as revealed by co-staining with the norepinephrine transporter (NET) (Figure S2). This anatomical convergence of noradrenergic input from the LC and inputs from the BLA onto MDm raised the possibility that norepinephrine (NE) may modulate the BLA→MDm synapse. Indeed, the BLA is heavily influenced by stress through stress-induced neuromodulation, and NE enhances BLA excitability and BLA-dependent behaviors such as fear learning (Arnsten, 2009; Likhtik and Johansen, 2019). To explore a possible role for NE in BLA→MDm transmission, we first performed immunostaining against the norepinephrine transporter (NET), known to label noradrenergic axons (Aggarwal and Mortensen, 2017; Pérez-Santos et al., 2021), in sagittal slices of MDm from mice that had been injected with AAV2/5-Syn-ChR2-mCherry into the BLA. These experiments revealed an abundance of NET+ fibers travelling throughout MDm, oftentimes in close proximity to mCherry+ BLA axons and puncta (Figure 4A, C). This observation encouraged us to examine the functional effect of NE on BLA→MDm synaptic transmission.

We performed similar whole-cell voltage clamp recordings as before (Figure 2A-G). Bath application of 10 µM NE rapidly inhibited BLA→MDm oEPSCs for both AMPAR- and NMDAR-mediated components (Figure 4B). Across multiple cells, NE significantly reduced AMPAR and NMDAR peak amplitudes (Figure 4D-E). Additionally, NE significantly increased the PPR)of BLA→MDm oEPSCs (250 ms ISI), suggesting that NE reduces the vesicular release probability of BLA axons in MDm via a presynaptic mechanism (Figure 4F-G).

We next asked which noradrenergic receptor is responsible for the inhibitory effects we observed. NE primarily signals through three classes of adrenergic receptors – α_1_, α₂, and β – of which only α₂ is known to signal via inhibitory Gi signaling (Perez, 2020). We wondered whether we could reverse NE’s inhibitory effects on BLA→MDm oEPSC amplitudes by subsequent bath application of yohimbine, an α₂-adrenergic receptor antagonist, in the presence of NE (Figure 5A, example currents and time course). Indeed, subsequent wash-in of yohimbine after NE-induced inhibition partially restored oEPSC amplitudes across multiple cells (Figure 5B-C). Additionally, yohimbine was able to partially restore the paired-pulse ratio of BLA→MDm oEPSCs to a similar level as before NE application (Figure 5D). Lastly, to further confirm the role of α₂-adrenergic receptors at the BLA→MDm synapse, we tested the effects of a selective agonist, as yohimbine may have off-target binding onto other receptors at high concentrations (MacDonald et al., 1997). Bath application of UK14304, a highly selective α₂-adrenergic receptor agonist with no known off-target binding to other receptors (MacDonald et al., 1997; Proudman et al., 2022), reproduced NE’s effects on BLA→MDm AMPAR and NMDAR oEPSC amplitudes (Figure 6A-C). UK14304 also increased the PPR of these synapses (Figure 6D), to a similar degree as NE.

**Figure 5.**
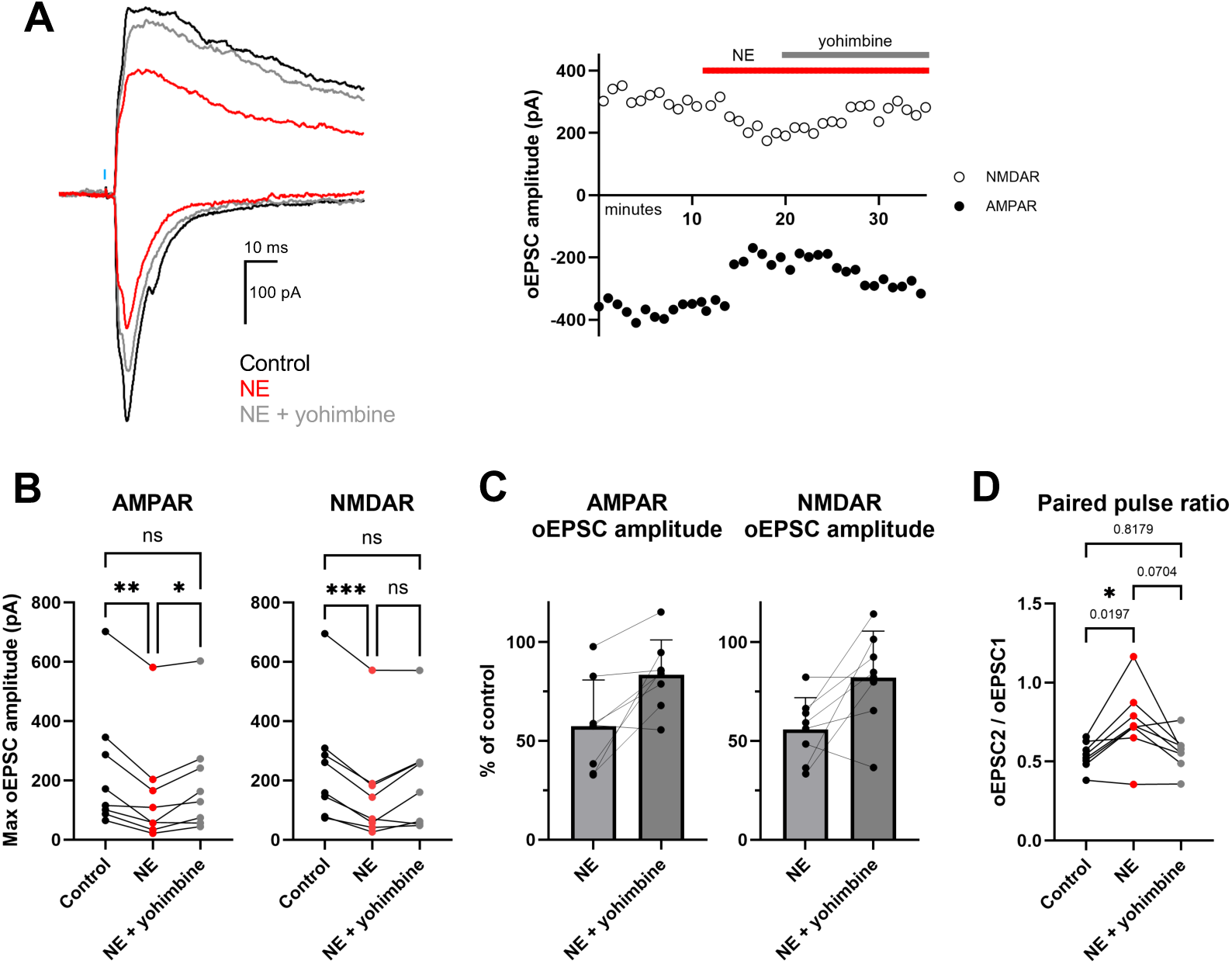
NE acts in part through α₂-adrenergic receptors to modulate the BLA→ MDm synapse. ***A***, Example oEPSCs from BLA→MDm inputs. Yohimbine, an α₂-adrenergic receptor antagonist, partially reversed NE-induced reduction of oEPSC amplitudes when applied together with NE. Black = control, red = 10 μM NE application, grey = simultaneous 10 μM NE and 20 μM yohimbine application. Blue bar indicates time point of 0.2 ms light stimulation. Right, time course of oEPSC peak amplitude during the experiment. Black filled circles = AMPAR, white circles = NMDAR. ***B***, Summary of effects of NE application on peak AMPAR (F(2,7) = 15.18, p = 0.0008) and NMDAR (F(2,7) = 17.28, p=0.0004) oEPSC amplitudes. Yohimbinepartially reversed the effect. ***C***, oEPSC amplitudes from ***B*** normalized as a % of pre-drug control. Normalized AMPAR and NMDAR oEPSC amplitudes during the NE + yohimbine condition (AMPAR 83.47% ± 17.65% and NMDAR 82.09% ± 22.55%) were larger than during NE-only application (AMPAR 57.47% ± 23.34% and NMDAR 55.82% ± 16.16%). ***D***, There was also a significant effect of drug condition on BLA→MDm oEPSC paired pulse ratio (PPR) (F(2,7) = 8.172, p = 0.0098). PPR values for control, NE, and NE + yohimbine conditions were 0.54 ± 0.09, 0.75 ± 0.23, and 0.56 ± 0.12, respectively. *Statistics* (***B****, **D***): One-way repeated measures analysis of variance (ANOVA). Post-hoc Tukey tests with corrections for multiple comparisons. (***B – D***): Values are mean ± SD. N = 8 cells / 7 mice. Lines connect values from the same cell.

**Figure 6.**
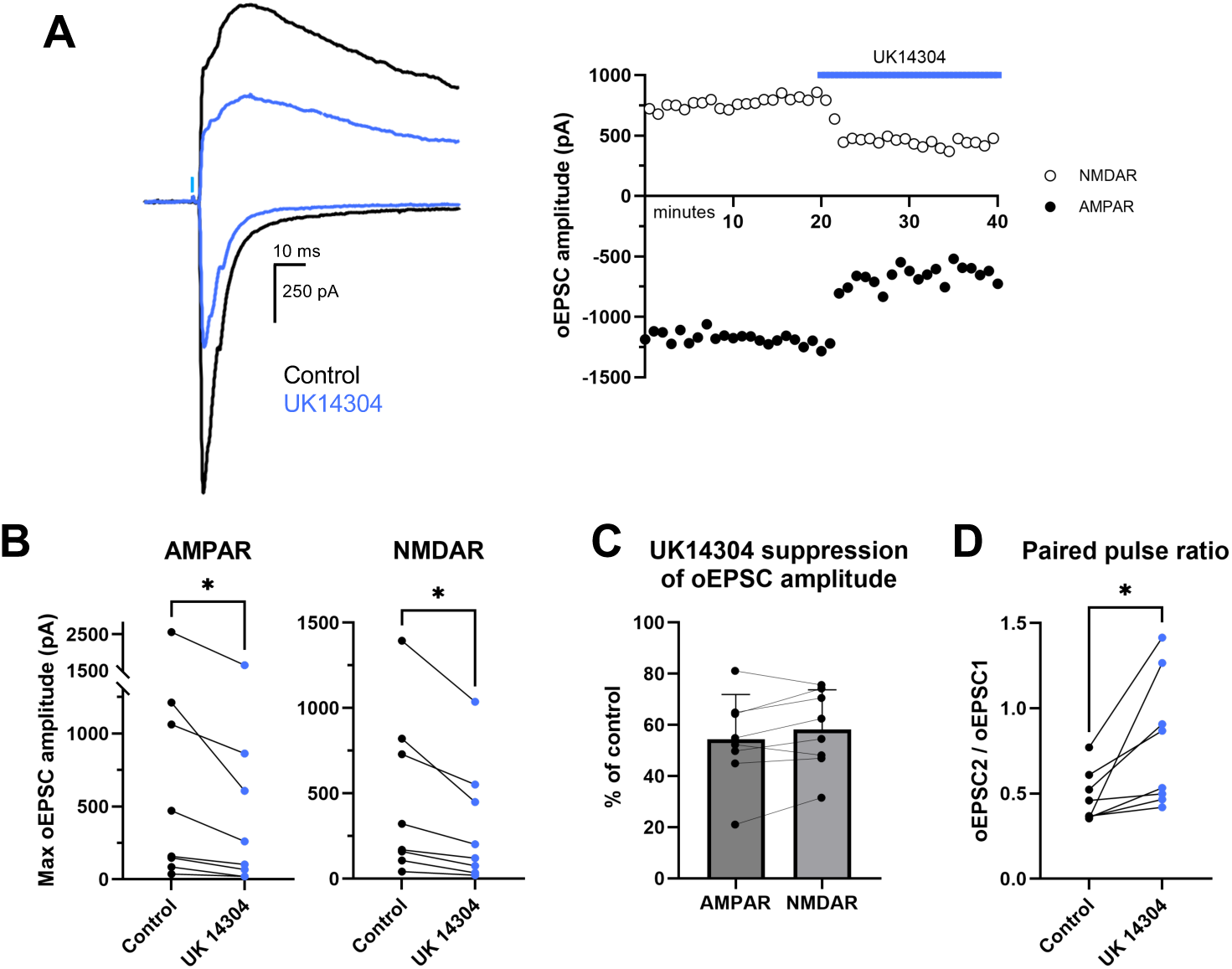
UK14304, a specific α₂-adrenergic receptor agonist, reproduces the effect of NE on BLA→ MDm oEPSCs. ***A***, Example recording of BLA→MDm oEPSCs, before and during bath application of 20 μM UK14304. UK14304 reduced the amplitudes of both AMPAR and NMDAR currents. Black = control, blue = UK14304 application. Light blue bar indicates time point of 0.2 ms light stimulation. Right, time course of oEPSC peak amplitude during the experiment. Black circles = AMPAR, white circles = NMDAR. ***B***, UK14304 significantly reduced the peak amplitudes of AMPAR (t_(7)_=2.370, p=0.0496) and NMDAR (t_(7)_=3.255, p=0.0140) mediated oEPSCs. ***C***, Plot of oEPSC amplitudes from ***B*** during UK14304 application normalized as a % of pre-drug control. UK14304 application significantly reduced AMPAR and NMDAR oEPSCs to 54.32% ± 17.52% and 61.67% ± 13.13% of their original pre-drug amplitude. ***D***, Summary of the effects of UK14304 on short-term plasticity. UK14304 significantly increased the paired pulse ratio of BLA→MDm AMPAR oEPSCs from 0.48 ±0.15 to 0.80 ± 0.38, t_(7)_ = 2.902, p=0.0229. *Statistics (**B, D**)*: Pairwise t-tests. (***B - D***): Values are mean ± SD. N = 8 cells/8 mice. Lines connect values from the same cell.

**Figure 7.**
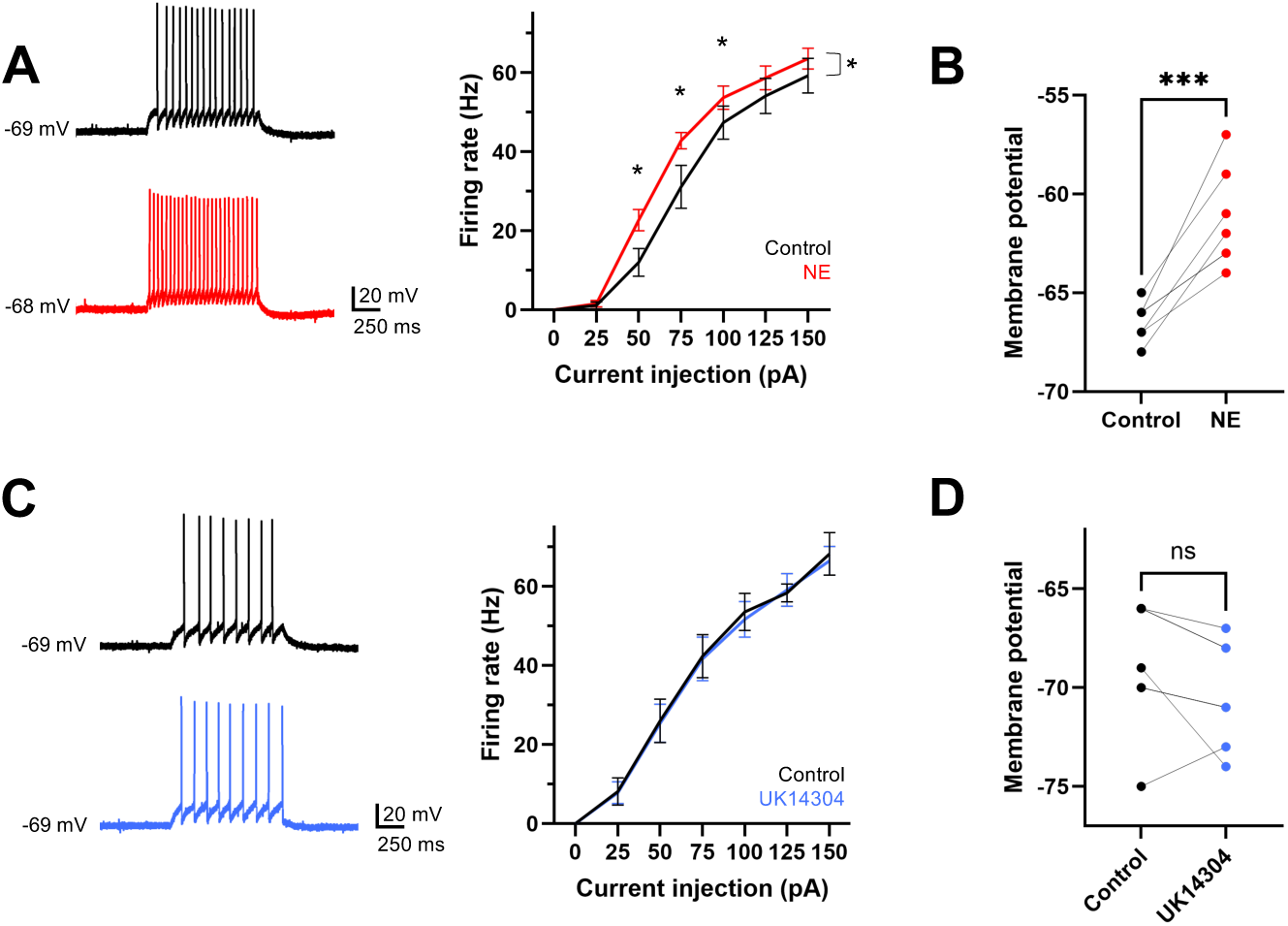
Norepinephrine, but not UK14304, increases the intrinsic excitability of MDm neurons. ***A-B***, Whole cell current clamp experiments were performed with MDm neurons while injecting holding current to maintain a resting membrane potential of -67 ± 3 mV. One second current steps were applied from 0 to +150 pA in +25 pA intervals to examine spiking responses in the presence of synaptic blockers. ***A***, Left, example neuron’s response to one second of 50 pA injection; before (top) and during (bottom) bath application of NE. Right, graph of firing rate with respect to current injection for control (black) and during bath application of NE (red). Values plotted are mean ± SEM. NE significantly increased the firing rates of MDm neurons (F(1,6) = 11.67, p = 0.0142, repeated measures 2-way ANOVA) with a significant interaction between drug condition and current injected, F(6,36) = 4.356, p = 0.0021. N = 7 cells / 3 mice. ***B***, Same neurons from ***A*** before applying holding current. NE significantly increased the resting membrane potential of MDm neurons from -66.4 ± 1.0 mV to -61.3 ± 2.5 mV for an average increase of 5.1 ± 2.3 mV (t_(6)_=6.00, p=0.001, paired t-test, mean ± SD. ***C***, Left, example of an individual neuron’s response to one second of 50 pA injection; before (black) and during (blue) bath application of UK14304, respectively. Right, graph of firing rate with respect to current injection for control and during bath application of UK14304. Values plotted are mean ± SEM. UK14304 did not significantly change MDm firing rates in response to depolarizing current injections (F(1,6) = 0.2289, p = 0.6493, mixed-effects ANOVA), N = 7 cells / 3 mice. ***D***, Same neurons from ***C*** before applying holding current. Some neurons showed a slight hyperpolarization in membrane potential from -68.9 ± 3.3 mV to -70.3 ± 2.7 mV after UK14304 application, though these changes were not statistically significant (t_(6)_=1.826, p=0.1177, paired t-test). Values are mean ± SD.

### Noradrenergic modulation of MDm excitability

Our findings are consistent with a role for NE in suppressing BLA→MDm transmission through α₂-adrenergic receptors and reducing the release probability of BLA axon terminals. To determine whether NE also exerts postsynaptic effects, we examined possible effects of NE on MDm neuron excitability. We performed current clamp experiments on MDm neurons in the presence of synaptic blockers and recorded MDm neuronal firing rates in response to a range of current injections.

We found that NE significantly increased MDm firing rates in the tonic mode (Figure 7A). NE also induced a significant depolarization of MDm neurons (Figure 7B). In contrast, UK14304 did not significantly change MDm neuron firing rates or membrane potentials in the tonic mode (Figure 7C-D). Additionally, NE and UK14304 did not significantly change the number of burst spikes that MDm neurons fired after release from hyperpolarizing current injections (Figure S5). Thus, NE depolarizes MDm neurons and increases their intrinsic excitability, increasing their responses to excitatory inputs while in tonic mode.

## DISCUSSION

### BLA sends a strong net excitatory input to MDm

Here, we further characterized the synaptic connection between BLA and MDm. We first confirmed that the BLA sends an anatomically restricted input to MDm in the mouse as previously described (McDonald, 1987; Hintiryan et al., 2021). We then confirmed that BLA forms glutamatergic inputs onto MDm neurons (Ahmed and Paré, 2023) that are mediated by AMPA and NMDA receptors. We also found that BLA→MDm oEPSCs exhibit paired-pulse depression, indicating a high probability of release. Additionally, the amplitude of single fiber oEPSCs were a fraction of maximal BLA→MDm oEPSCs, suggesting that multiple BLA inputs can converge onto a single MDm neuron. Some of these inputs are strong enough to evoke MDm neuron spiking alone or in combination with the activation of other BLA inputs. The latter scenario is particularly interesting given that BLA neurons are heterogeneous in their responsivity to rewarding vs. punishing stimuli (Janak and Tye, 2015; Kim et al., 2016; Lutas et al., 2019). Such convergence could explain the wide range in amplitudes of BLA→MDm maximal oEPSCs. Overall, our results suggest that the BLA sends key affective information, potentially of different types, to activate a subset of MDm neurons which, in turn, relay the information to downstream areas such as the mPFC (Anastasiades et al., 2021).

Our *ex vivo* sagittal slice preparation greatly facilitated the study of the BLA→MDm synapse. Prior studies have suggested the BLA→MDm synapse is weak or non-functional in the rodent (Máytás et al., 2014; Ahmed and Paré, 2023). Our initial pilot studies using coronal slices found that BLA→MDm oEPSCs were observed only infrequently (the incidence of any response was <33%, not shown). This motivated us to create a slice preparation preserving the BLA inputs to MDm to more reliably and accurately study this synapse. Our sagittal slice preparation improved our ability to identify MDm neurons postsynaptic to BLA, as the preservation of BLA axons travelling from the ventral amygdalofugal tract provides a visual aid for identifying potentially innervated MDm neurons (incidence of any response ∼64%; see example, Figure 1E). Additionally, we observed that processes of patched and filled MDm neurons were in the same plane and orientation as some BLA axons travelling in the anterior to posterior direction (example; Figure 4C), suggesting that we may have preserved more of the BLA→MDm connection with sagittal slices that otherwise could have been severed in the coronal orientation. These factors may contribute to an improved ability to investigate MDm neurons that receive strong BLA-induced excitatory drive. Overall, the preservation of the ventral amygdalofugal tract in our *ex vivo* slice preparation provides a method for electrical or optical stimulation of amygdalar inputs to MDm, setting the stage for future studies.

The BLA neurons that send direct inputs to mPFC (BLA→mPFC) are likely to be a separate subpopulation compared to those that send inputs to MDm (BLA→MDm) (McDonald, 1987). This raises the hypothesis that BLA→mPFC neurons may be carrying different types of information compared to BLA→MDm neurons. Indeed, BLA contains subpopulations of neurons that modulate either fear learning or fear extinction (Herry et al., 2008; Kim et al., 2016; Hochgerner et al., 2023). Interestingly, MD is *not* necessary for fear learning but is necessary for fear extinction; tonic spiking in MD enhances fear extinction, while burst spiking impairs it (Lee et al., 2012). Additionally, MD has been implicated in modulating cognitive and behavioral flexibility (Rikhye et al., 2018; Zhang et al., 2025), whereas damage to MD in humans induces attention and memory deficits, learning difficulties, and emotional instability (Zoppelt et al., 2003; Van Der Werf et al., 2003). Lesions of MDm also cause impairments in learning new associations while leaving previously learned associations intact (Mitchell and Chakraborty, 2013; Mitchell, 2015). Given the vital role of BLA and mPFC in fear extinction (Tovote et al., 2015), we speculate that input from BLA→MDm and onwards to mPFC (Georgescu et al., 2020; Anastasiades et al., 2021) may support this process. Additionally, the key role of BLA in emotional learning suggests that its input to MDm, and subsequent transmission from MDm→mPFC, support learning of new associations that contribute to behavioral flexibility and emotional stability.

### Noradrenergic modulation of BLA**→**MDm input

In addition to characterizing the glutamatergic input to MDm from BLA, we identified a novel neuromodulatory mechanism that robustly inhibits this input. Specifically, we showed that NE binds to α₂-adrenergic receptors to reduce synaptic transmission and lower the vesicular release probability of BLA→MDm inputs. Our results reveal a rapid and dynamic mechanism to transform affective information from BLA before it is relayed downstream in a behavioral state-dependent manner.

The inhibitory presynaptic actions of NE on BLA→MDm inputs stand in contrast to the role of NE in *enhancing* the excitability of BLA neurons (Likhtik and Johansen, 2019). Acute stress-induced NE release in BLA triggers persistent anxiety states to facilitate sensory perception, emotional memory consolidation, and anxiety-related behaviors conducive to survival during dangerous situations (McGaugh, 2004; Arnsten, 2009; Daviu et al., 2019). Indeed, NE release from LC activates β-adrenergic receptors in BLA to produce conditioned place aversion and anxiety-like behaviors (McCall et al., 2017). NE’s enhancement of BLA excitability *increases* BLA output to downstream regions such as the central amygdala (CEA) and ventral hippocampus (vHPC), resulting in enhanced anxiety and emotional memory consolidation (Felix-Ortiz et al., 2013; Namburi et al., 2015; Atucha et al., 2017; Beyeler et al., 2018). In summary, NE enhances BLA excitability and increases functional transmission from BLA to other regions, contributing to stress-related changes in cognitive processing and behavior. In light of this work, our finding that NE inhibits transmission from BLA→MDm highlights a potential state-dependent rerouting of information transmission from BLA to different targets.

What is the role of noradrenergic inhibition of BLA→MDm transmission? Arousal and stress-induced release of NE in BLA and mPFC impairs extinction learning (Giustino and Maren, 2018; Likhtik and Johansen, 2019; Giustino et al., 2020). Simultaneously, stress-induced NE impairs mPFC-dependent functions such as cognition, decision making, and behavioral flexibility (Arnsten et al., 2012). NE LC neurons are also activated by motivationally salient and/or unexpected stimuli and play a key role in learning and decision making (Poe et al., 2020; Breton-Provencher et al., 2021; Jordan and Keller, 2023; Basu et al., 2024). We hypothesize that an additional mechanism underlying stress-induced impairment of extinction learning is the inhibition of BLA→MDm transmission via stress-induced NE release into MDm. If BLA→MDm transmission supports learning and behavioral flexibility via downstream recruitment of mPFC, stress-induced elevations in NE could dampen this recruitment of mPFC and impair mPFC-dependent functions. These hypotheses are consistent with the role of stress in switching processing modes from “top-down” to “bottom-up” via enhancing amygdala-dependent functions and dampening cortical functions (Arnsten, 2009; LeDuke et al., 2023). Thus, MDm is a powerful relay between BLA and mPFC that is uniquely positioned to facilitate an NE-induced switch in cognitive processing.

NE signals through a diverse set of receptors, each with different binding affinities. α₂- adrenergic receptors have the highest affinity for NE (K_d_ ≈ 56 nM), with α_1_ (330 nM) and β subtypes (740 nM) having lower affinities (Ramos and Arnsten, 2007). Here, we have shown that NE-induced inhibition of excitatory BLA→MDm input is dependent upon α_2_-adrenergic receptors, while NE’s effects on enhancing MDm excitability are dependent upon adrenergic receptors *other* than α_2_ (such as α_1_ or β-adrenergic receptors). Thus, different types of adrenergic receptors may be activated depending on the amount of NE released by LC inputs to MDm, leading to diverse behavioral state and NE concentration-dependent effects on amygdalo-thalamic circuitry. Indeed, NE release by LC is directly proportional to arousal and stress levels (Giustino et al., 2020; Poe et al., 2020). During states with low levels of NE release, such as during low to moderate arousal, BLA→MDm input may be suppressed via activation of high-affinity α_2_-adrenergic receptors. As NE levels increase during states of high arousal and stress, MDm neurons may also become more excitable, thereby increasing the signal to noise ratio of strong BLA→MDm input, which is then relayed downstream while simultaneously enhancing transmission of information from other areas of the brain. Since action potentials are an all-or-nothing phenomenon, we hypothesize that NE may have the strongest silencing effects on BLA inputs that are more moderate in strength and closer to the rheobase of MDm neurons, as these inputs could become sub-threshold post-NE inhibition. On the other hand, very strong net input from BLA may still be able to evoke spiking post-NE inhibition, particularly if MDm neurons are at more depolarized potentials. Furthermore, as BLA shows high-frequency gamma oscillations during emotional memory consolidation (Kanta et al., 2019), NE could act as a high-pass filter during these processes by reducing the paired-pulse depression of the BLA→MDm synapse as previously described for other neuromodulators (Seeburg et al., 2004), thereby facilitating transmission of only the most salient information from BLA→MDm.

Here, we have shown that BLA sends a strong excitatory input to MDm that can evoke spikes in both tonic and burst modes. NE significantly reduces transmission at this synapse via α_2_-adrenergic receptors while enhancing MDm excitability via other adrenergic receptors (Figure 8). Taken together, these results suggest that NE dampens the net disynaptic relay from BLA→MDm→mPFC, in contrast to the NE-induced enhancement of BLA outputs to other brain areas (Daviu et al., 2019; Likhtik and Johansen, 2019). Thus, our study uncovers novel neuromodulatory mechanisms that are likely to modulate associative learning and decision making and provides insight into how stress changes amygdalo-thalamo-cortical circuitry to affect learning and behavioral flexibility.

**Figure 8.**
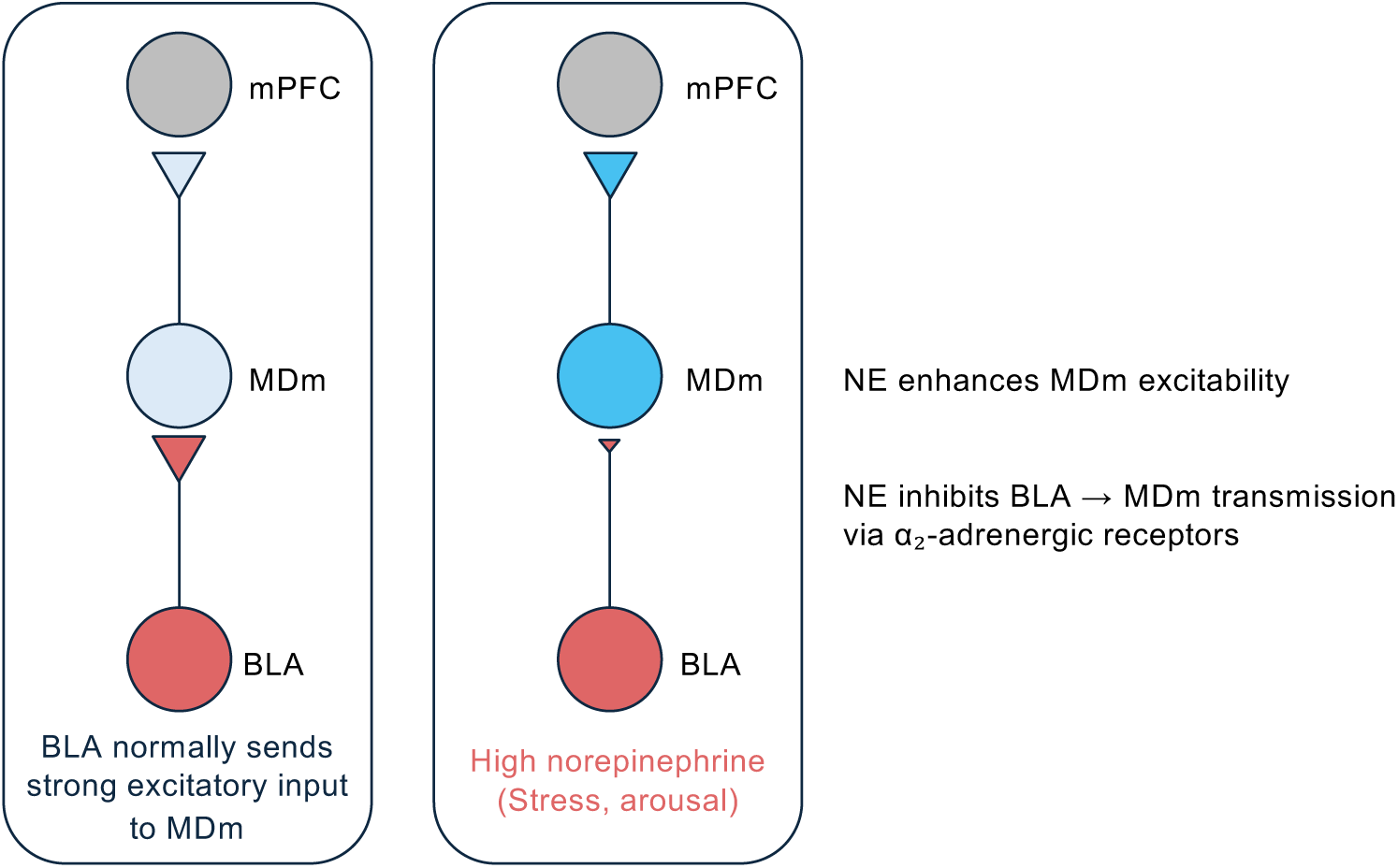
Model schematic of the BLA→ MDm→ mPFC circuit during different NE states. ***Left***, BLA sends affective information to MDm which is potentially further relayed from MDm to mPFC. ***Right***, model schematic of BLA→MDm→mPFC circuit during high NE states, which occur during times of high physiological stress or arousal. BLA→MDm synaptic transmission is significantly inhibited by NE binding onto α₂-adrenergic receptors, which reduces the vesicular release probability of BLA axonterminals. Simultaneously, high levels of NE increase MDm intrinsic excitability, increasing the ability of other inputs to evoke MDm spiking while information flow via BLA input is reduced. Thus, NE may shift information flow such that MDm activity is more heavily driven by other inputs from other areas of the brain instead of from the BLA, changing the nature of the information that MDm sends to mPFC.

## Acknowledgements

We thank members of the Chen and Andermann labs for helpful discussions about the project and manuscript. Support was provided by the National Institute of Health (NIH) R01EY032749 to C.C. and M.L.A., NIH R01MH123431 to M.L.A., NIH R01EY013613 to C.C., and Tan-Yang Center for Autism Research Grant to C.C. We would like to thank the Administrative and BCH Cellular Imaging Core (RRID:SCR_026485) funded by NIH P50 HD105351, S10OD016453 and NIH S10OD030322, and the Viral Core funded by NIH P30EY012196.

