## Supplemental Figures for "Noradrenergic Modulation of an Amygdalo-thalamic Circuit"

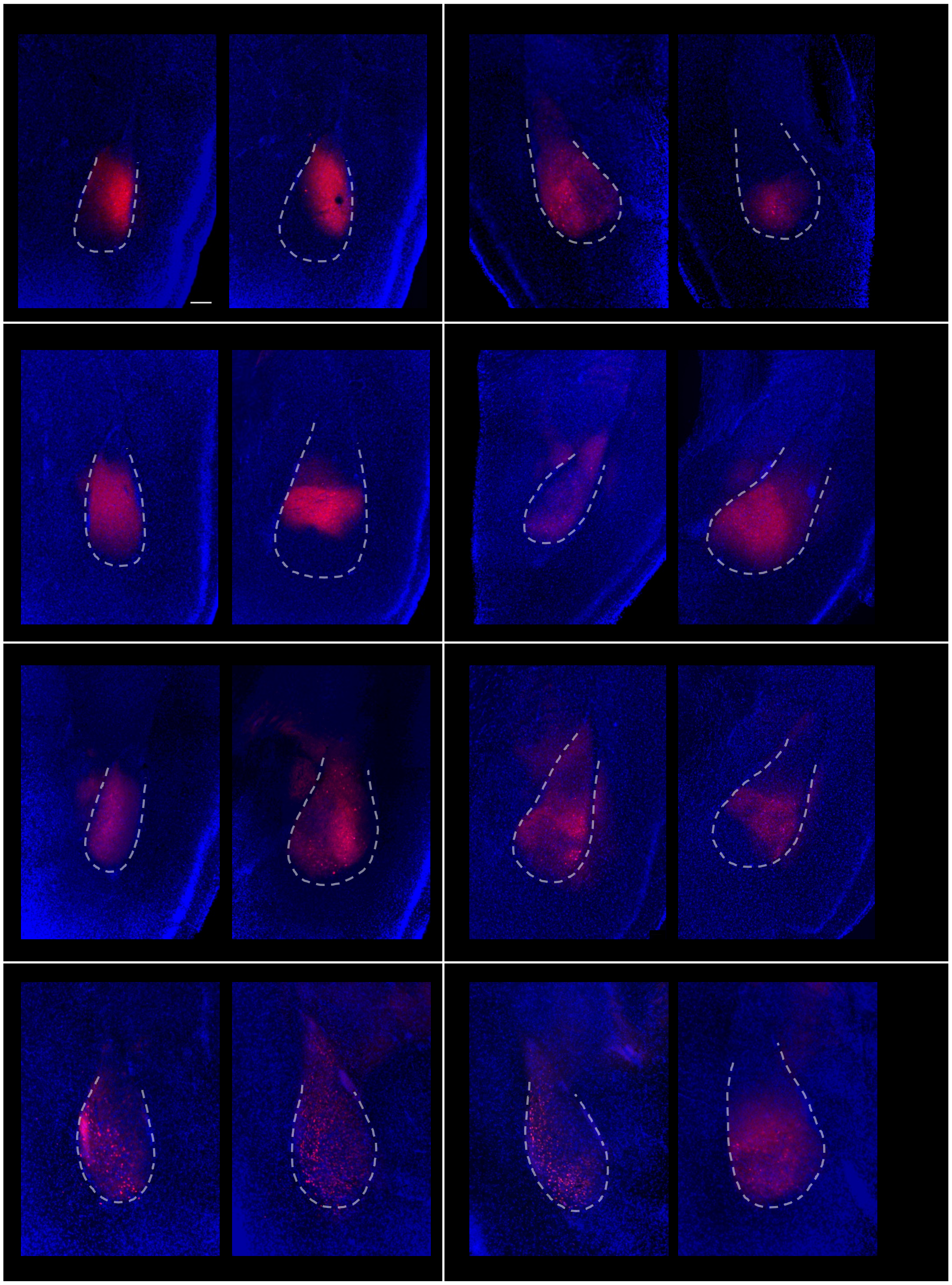

**Supplemental Figure 1A. Representative examples of stereotaxic injections into BLA**
***A*** Images from coronal sections of BLA following injection of AAV2/5 Syn-ChR2-mCherry from 8 representative mice from which we recorded BLA-> MDm synaptic currents. For each mouse (grid), pairs of images from the anterior (left) and posterior (right) aspects of the BLA is shown. White dotted line outlines the perimeter of BLA based on the location of the amygdalar capsule and the density of the DAPI stained nuclei.

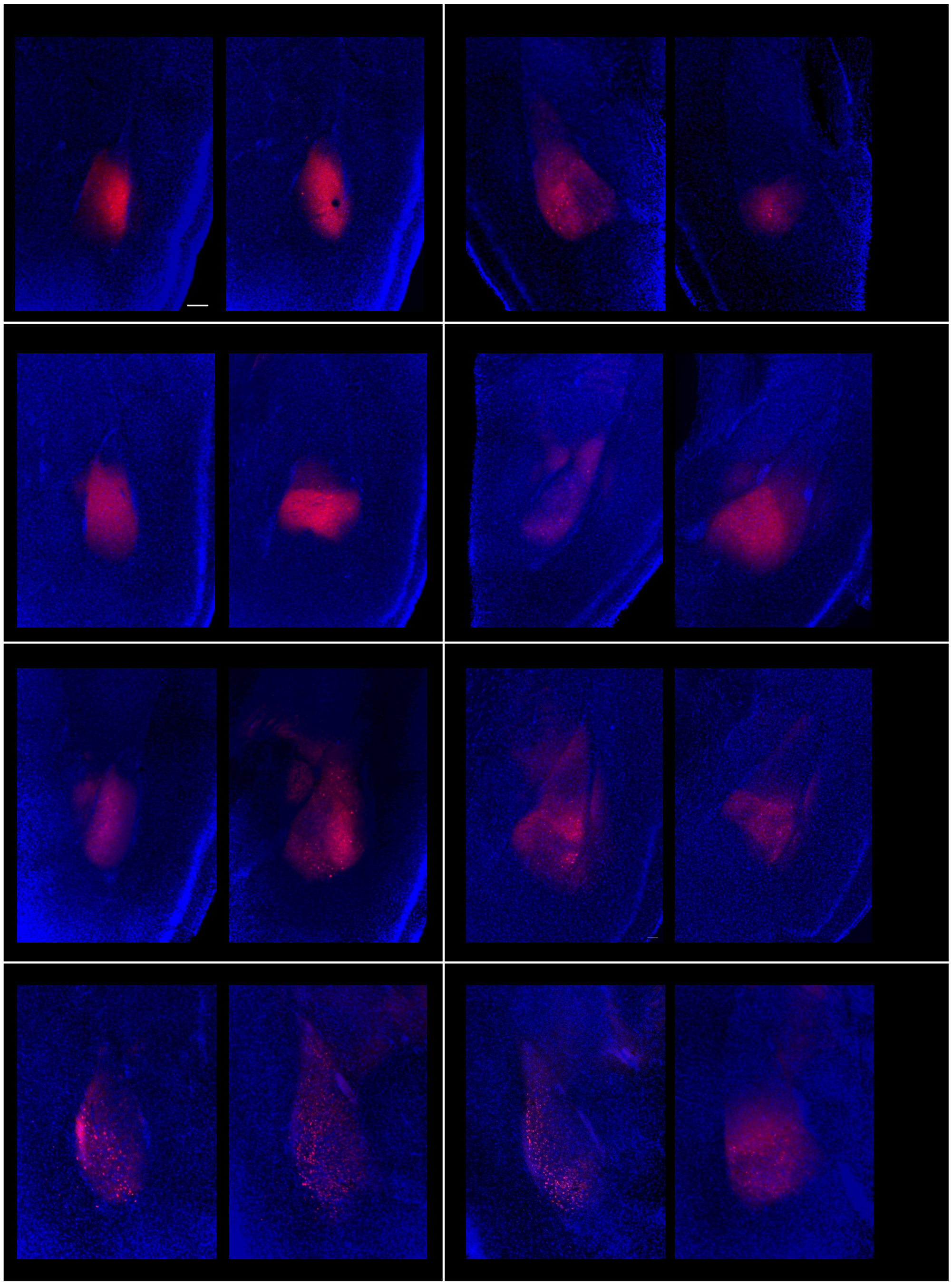

**Supplemental Figure 1B. Representative examples of stereotaxic injections into BLA (continued)**
***B*** the same images as in A but without the white dotted line. Scale bar=100 µm.

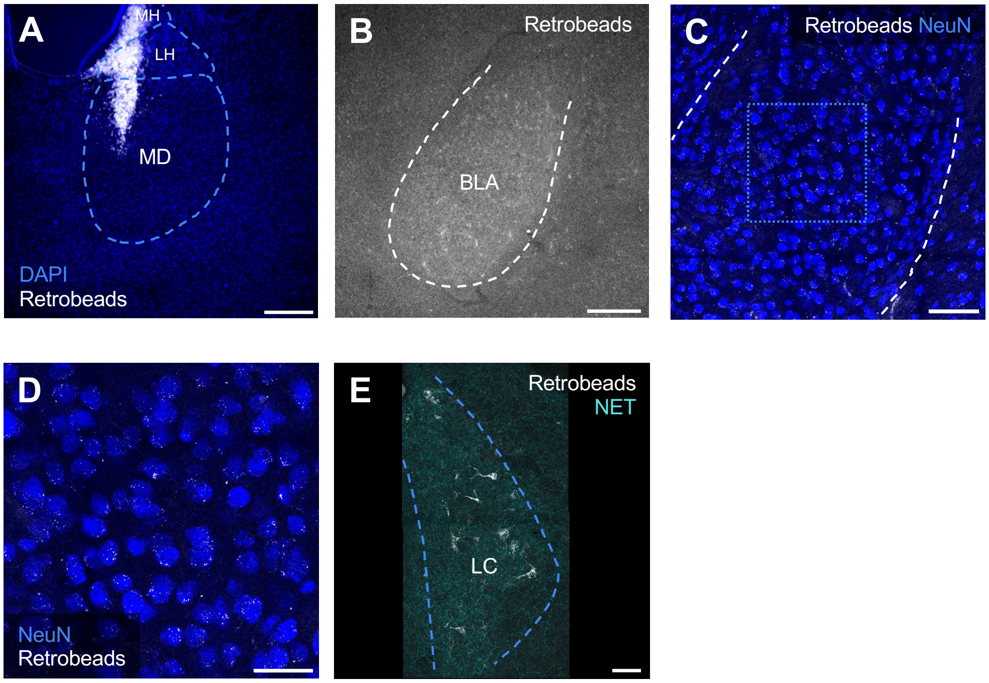


**Supplemental Figure 2. Locus coeruleus sends inputs to MDm.**
***A***, Retrobeads were injected into MDm where BLA axons terminate as shown in Figure 1B. Scale bar = 200 µm. ***B***, Image shows retrobead labeling in BLA, shown at a lower magnification level. Scale bar = 200 µm. ***C,*** BLA from ***B*** at a higher magnification to allow visualization of retrobeads (white). BLA neurons were stained with NeuN (blue). Scale bar = 100 µm. ***D,*** Higher magnification of BLA area outlined by the dashed square in ***C***. Scale bar = 50 µm. ***E***, Image of the ipsilateral locus coeruleus (LC) which was retrogradely labeled by the MDm retrobeads injection shown in ***A***. Staining against NET was used to identify the LC. Scale bar = 50 µm.

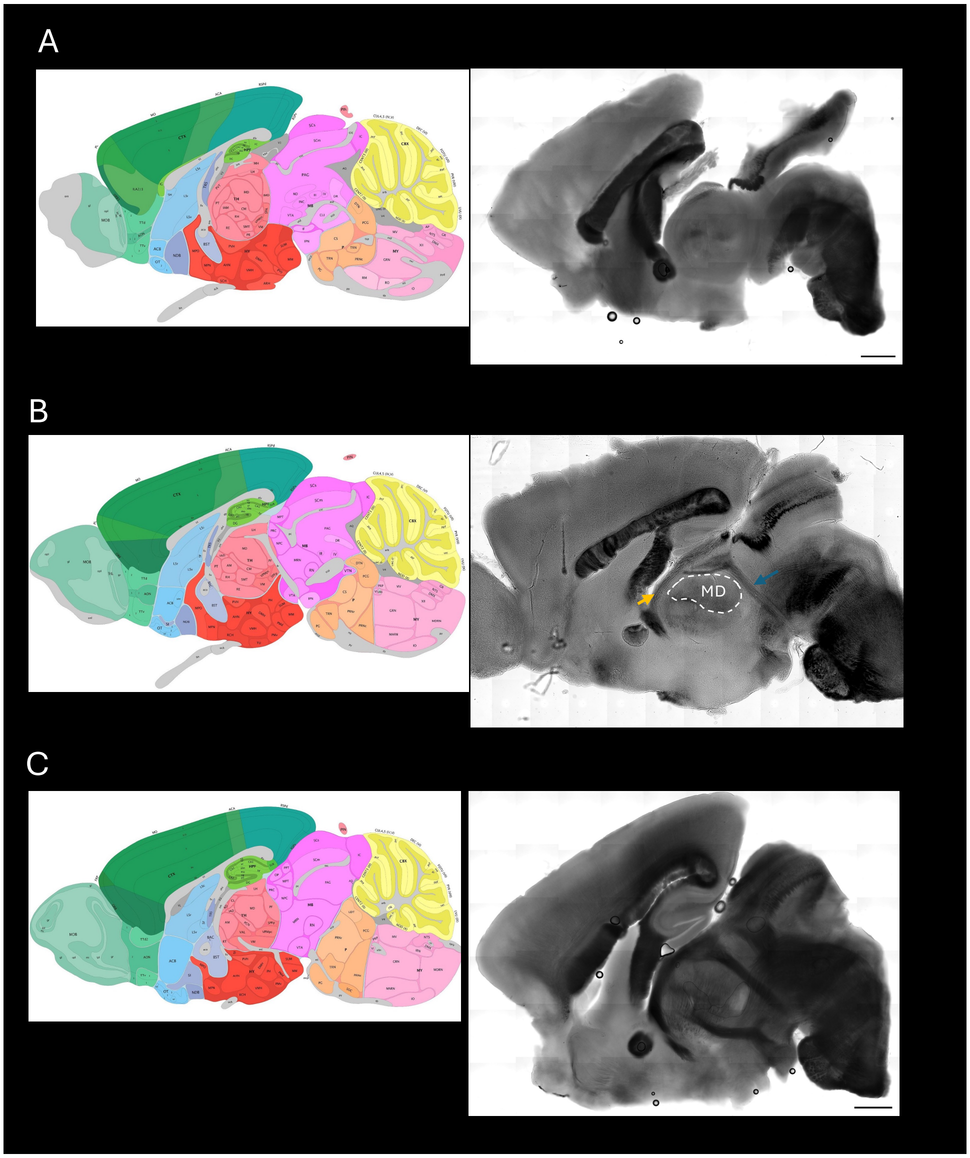


**Supplemental Figure 3: Saggital Slice Preparation of MDm**

Images from the Allen Brain Atlas (left) and images from comparable brightfield slices of adult mice (right) ordered from most medial section (A) to lateral section (C). We find the optimal slice to obtained intact BLA inputs onto MDm neurons is section (B). Landmarks of this section include the fiber tract containing the habenula-interpeduncular tract (indicated by blue arrow), and the fiber tract containing the ventral amygdalofugal tract (indicated by the orange arrow). We avoid Section (A) as it is too medial and contains cells from PVT juxtaposed to the 3rd ventricle. Connected cells are more difficult to find in section (C) where the MD nucleus is no longer juxtaposed to the fiber tract containing the ventral amygdalofugal tract.

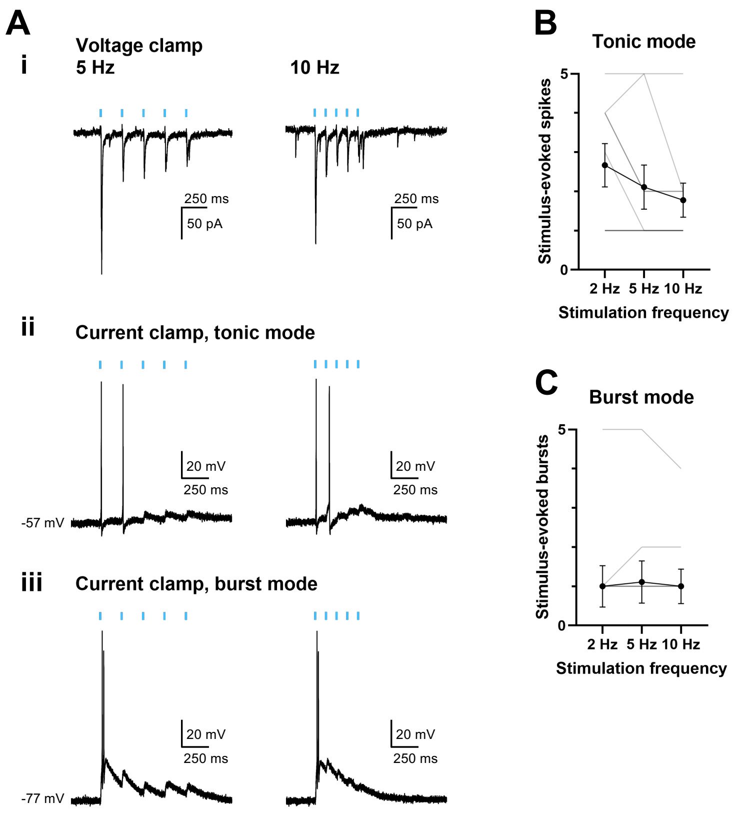


**Supplemental Figure 4. Optogenetic stimulation of BLA inputs at higher frequencies.**
Related to Figure 3. ***A***, BLA inputs were stimulated with 5 0.2-ms pulses of 10 $mW/mm^2$ 470 nm light (blue bars) at 5 Hz (left column) and 10 Hz (right column) while recording in voltage clamp (*i*), current clamp (tonic mode) (*ii*), and current clamp (burst mode) (*iii*) in the same MDm neuron. ***B – C,*** Black lines and error bars indicate mean +/- SEM, grey lines indicate individual neurons. N = 9 cells / 4 mice. ***B,*** Mean number of stimulus-evoked spikes for each stimulation frequency during tonic mode. ***C,*** Same cells from ***B***. Mean number of stimulus-evoked bursts for each stimulation frequency during burst mode.

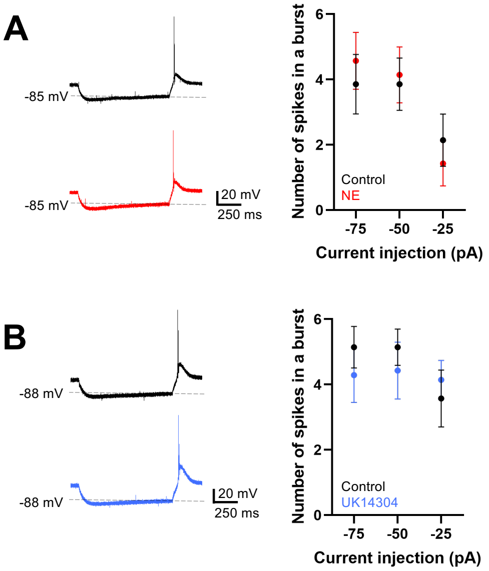


**Supplemental Figure 5. Norepinephrine and UK14304 do not significantly change MDm burst spiking.** ***A-B***, Same neurons and recording conditions as Figure 7. 1 second current steps were applied from -75 pA to 0 pA in +25 pA intervals. Cessation of hyperpolarizing current steps resulted in rebound burst spiking which was quantified before and during bath application of 10 µM NE or 20 µM UK14304. ***A***, Left, example of a neuron's response to one second of -50 pA injection before and during bath application of NE. Right, graph of spikes in a burst with respect to hyperpolarizing current injection, before (black) and during NE application (red). Values plotted are mean ± SEM. NE did not significantly affect burst spiking ($F(1,6) = 0.1600$, $p = 0.7030$, repeated measures 2-way ANOVA). N = 7 cells / 3 mice. ***B,*** Left, example of a neuron's response to one second of -50 pA injection; top and bottom before (black) and during UK14304 application (blue). Right, graph of spikes in a burst with respect to hyperpolarizing current injection. Values plotted are mean ± SEM. UK14304 did not significantly affect burst spiking ($F(1,6) = 0.2530$, $p = 0.6329$, repeated measures 2-way ANOVA). N = 7 cells / 3 mice.
